# Mutualistic microbes drive niche expansion yet constrain diversification

**DOI:** 10.64898/2026.01.14.699472

**Authors:** Eleanor M. Hay, Rosana Zenil-Ferguson, Michelle E. Afkhami, Christopher A. Searcy

## Abstract

Mutualistic interactions are widely hypothesized to expand ecological niches and promote diversification, yet evidence for this relationship at macroevolutionary scales remains limited. Here, we investigate how fungal seed endophytes (*Epichloë*) influence niche breadth and diversification across over 100 species of the globally-distributed grass genus *Poa*. Combining global-scale climate, environmental, phylogenetic, and symbiosis data, we constructed species distribution models for suites of symbiotic and non-symbiotic plant species, then used a range of phylogenetic comparative methods to assess the influence of these symbionts on niche breadth, niche breadth evolution, and diversification. Endophyte-positive species had ∼1.4 times broader primary niche axes (most often associated with temperature tolerances), ∼2.5 times larger geographic ranges, and >30 times faster rates of niche breadth evolution than non-symbiotic species. Endophytic plants also had significantly drier climate centroids, consistent with endophyte-mediated drought tolerance, while simultaneously showing constraints along specific niche dimensions. Despite pronounced niche expansion, hidden state speciation and extinction models revealed two modes of diversification: endophytes decrease diversification rates in young lineages, and have a neutral effect in older lineages. Together, our results demonstrate that mutualists can enhance ecological flexibility while simultaneously constraining macroevolutionary outcomes, revealing a decoupling of niche expansion and diversification, and highlighting the complex role of symbiosis in shaping host evolutionary trajectories.

## INTRODUCTION

Mutualistic relationships, in which both partners benefit^1^, are ubiquitous across the Tree of Life and can profoundly influence the ecology and evolution of their hosts. Mutualists provide a range of benefits, including defense against enemies^2^, enhanced nutrient acquisition^3^, and tolerance to environmental stress or pathogens^4^. These interactions often directly benefit host fitness by increasing growth, survival, and reproduction. For example, alfalfa plants associating with rhizobia show increased growth and fitness^5^, corals that host amphipods benefit from greater survival and growth^6^, and plants with diverse pollinators have enhanced reproductive success^7^. At the same time, the benefits provided by mutualists are complex and often context dependent, and can vary with host genotype, environmental conditions, and the presence of other symbionts^8,9^. A central unresolved question is whether the fitness benefits of mutualisms scale beyond individual interactions to influence niche evolution, diversification, and long-term macroevolutionary trajectories of host lineages.

Over evolutionary timescales, the benefits conferred by mutualists can extend beyond individual fitness to influence host lineages at broader scales. Mutualistic associations have the potential to facilitate colonization of new environments, drive expansion of broader ecological niches, and promote diversification. By relaxing ecological constraints, mutualists can enable persistence under otherwise unfavorable conditions, which can translate to broader niches and expanded range limits. For example, partnering with mycorrhizal fungi enhances plants’ access to soil nutrients and water, and enables them to tolerate a wider range of environmental conditions^10,11^. Similarly, partnering with ants, which provide protection from predation, allows butterflies to expand their niche to include novel plant hosts^12^. These benefits are also expected to influence diversification: larger geographic ranges are understood to buffer against extinction^13^ and facilitate speciation^14^, and expansion into novel niches is a key driver of evolutionary diversification and adaptive radiations^15^. Empirical studies support these ideas: frugivory in primates is associated with larger geographic ranges and increased diversification^16^, and symbiotic gall-inducing insects have broader ranges of host plants and are >17 times more diverse than their non-symbiotic relatives^17^.

Similar to processes occurring at the individual-level, the net macroevolutionary outcome of mutualism is highly context-dependent, shaped by variation in partner specificity, the level of partner dependence, and environmental setting^18^. Such factors could explain why the empirical effects of mutualisms on diversification remain conflicting: mutualists can have positive, neutral, or even negative effects on diversification^19^. Despite the benefits offered by mutualists, they can also impose evolutionary constraints, and co-dependence may stabilize niches, limit host distributions, or increase extinction risk if a partner is lost^20–22^. Because the same processes that affect diversification are also linked to ecological opportunity and environmental tolerance, understanding how mutualists influence niche breadth is essential to understanding mutualism’s role in evolutionary trajectories. Despite this connection, few studies have simultaneously examined how mutualists influence both niche breadth and diversification. Addressing this gap is key to linking individual-level mutualist effects to large-scale patterns of biodiversity.

Grasses (Poaceae) and their associations with clavicipitaceous fungal endophytes provide a powerful system for testing how mutualists impact niche breadth evolution and diversification. With over 12,000 species, grasses are one of the most important plant lineages on earth, and it is estimated that 20 – 30% of grass species have symbiotic relationships with fungal seed endophytes^23^. In particular, the heritable, fungal endophytes from the genus *Epichloë* associated with cool-season grasses (Pooideae subfamily) are mutualists that have the potential to alter their host’s evolution. *Epichloë* live intercellularly within host tissues, including reproductive structures, and are transmitted from parent to offspring via seeds^24^, with imperfect transmission acting as a mechanism by which endophytes can be lost^25^. Because their fitness depends on host reproduction, vertically transmitted endophytes are tightly aligned with host evolutionary dynamics^26,27^. These endophytes are known to provide a range of benefits to hosts such as conferring drought tolerance^28,29^, herbivore resistance via the production of alkaloids^30,31^, and other stress-buffering traits in exchange for photosynthetic carbon^32^. Benefits of associating with *Epichloë* endophytes include many outcomes beyond the level of individual fitness, including enhanced population growth^33^, range expansion^34^, and increased community-level dominance^35^. Yet, despite this extensive ecological research, we still know little about how these symbionts influence macroevolutionary processes like niche breadth evolution or diversification.

We focus on an ecologically and agriculturally important clade within cool-season grasses, *Poa* (Bluegrasses). Comprising over 600 species, *Poa* is the largest grass genus, spanning temperate regions worldwide, and has been recognized as an exceptional radiation^36^. Within *Poa*, past studies have suggested that *Epichloë* endophytes play a role in niche partitioning^37^, and can contribute to range expansions, such as the invasion of *Poa annua* into Antarctica^38^. We use *Poa* to test the hypothesis that mutualistic relationships with fungal seed endophytes increase host niche breadth (i.e., niche expansion) and drive host diversification. To achieve this, we combine global-scale host-endophyte data, species distribution modeling, and phylogenetic comparative analyses to evaluate how mutualistic fungal endophytes shape ecological niche breadth and diversification across an exceptional grass radiation.

## RESULTS

### Fungal endophytes enable their hosts to occupy drier climates

We determined which species of *Poa* are known to have associations with *Epichloë* endophytes by conducting a literature search and screening seeds from USDA germplasm banks. In total, we obtained information on endophyte status for 116 of the 606 recognized *Poa* species, representing ∼19% of the genus (Table S1-S2). This detailed survey revealed that 50 species of the evaluated *Poa* have been previously found to host endophytes, while another 66 evaluated species showed no evidence of hosting fungal seed endophytes. This ∼43% (95% CI: 34 – 52%) positive rate is slightly higher than early estimates of 20 – 30% for endophyte associations in grasses (Poaceae)^23^ and the proportion we found in manually screened seeds (7/37 species E+, 21.7%; Table S3). To combat possible publication bias for reporting endophyte-positive species, researchers were contacted directly when endophyte-negative taxa were not reported. We mapped 107 of 116 species to a previously published phylogeny^39^. The estimated crown age of *Poa* in this tree is 5.1 Mya, which falls within previous estimates for this clade^40,41^. We found that *Epichloë* endophytes are widespread across the *Poa* phylogeny (Figure 1a) and exhibit moderate phylogenetic signal with a D-statistic of 0.71. This was significantly different from both a random distribution (p = 0.013) and a Brownian motion model (p = 0.003) and indicates that endophytes are phylogenetically clustered across the tree, but less strongly than expected under a strict Brownian motion model.

**Figure 1.**
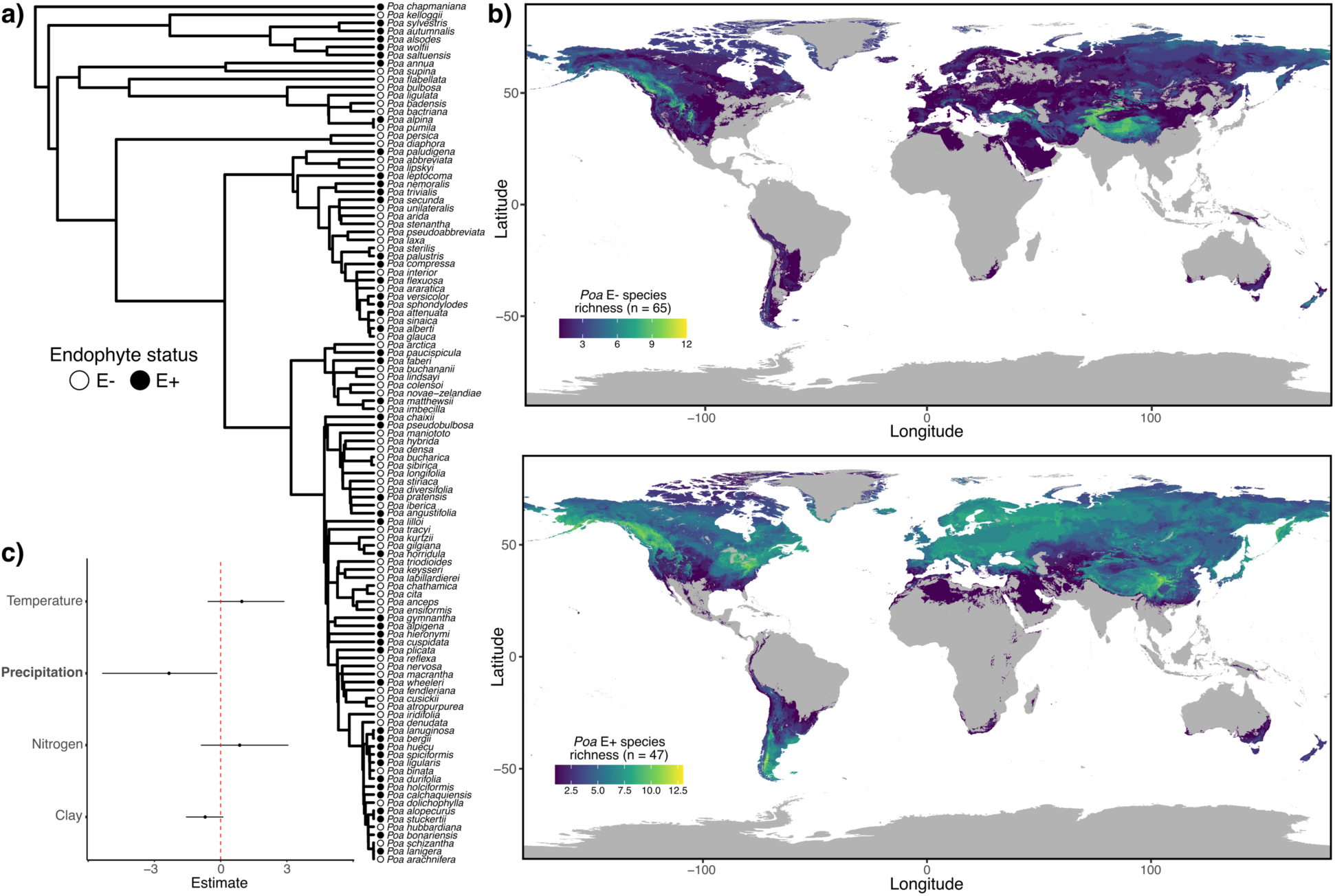
Phylogenetic and spatial patterns of *Epichloë* endophytes across *Poa.* A) displays the *Poa* phylogeny (n = 107) with dots at the tips indicating endophyte status: white = no known relationships with endophytes (n = 60); black = known associations with *Epichloë* endophytes (n = 47). B) shows spatial patterns of *Poa* species richness for endophytes-negative species (top) endophyte-positive species (bottom). C) displays mean and 95% confidence intervals of effects from the Bayesian spatiophylogenetic model testing the association between endophyte presence and key niche axes.

Niche breadth of *Poa* species was assessed using three metrics: geographic range size, range on primary niche axis, and niche hypervolume. Geographic range size and range on the primary niche axis were estimated using species distribution modeling^42^, and hypervolumes were calculated in multivariate environmental space^43^. The niche models performed relatively well for *Poa* species, with mean test AUC = 0.85 across species (Table S4; Figures S1-S12). This compares favorably with mean AUC scores produced by MaxEnt experts when modeling large suites of species (mean AUC = 0.59 – 0.81^44^). Of the 112 species that had enough occurrence records (>15) to construct niche models, 10 had models with test AUC < 0.7. The worst performing species was *Poa chathamica* (test AUC = 0.42), which is restricted to the Chatham Islands east of New Zealand. The remaining poorly performing models included all species with the greatest number of occurrence records (>10,000) and might reflect the difficulties in identifying niche limitations for these generalist species (Table S4).

To identify range on the primary niche axis, we extracted the top ranked climate/environmental variable from each species’ niche model and calculated the range on this axis. The most important variables identified in the niche models were largely temperature variables. Specifically, the top three variables were mean temperature of the warmest quarter (BIO10; top variable for 20 species), mean annual temperature (BIO1; top variable for 13 species), and temperature seasonality (BIO4; top variable for 12 species; Table S4). Given that *Poa* are temperate grasses, and are not found in tropical areas, it is unsurprising that temperature plays a primary role in determining species range limits.

Endophyte-positive species are more widespread and show greater range overlap than E-species, despite there being more *Poa* species without endophytes. Notably, mean range overlap among E+ species (measured as the Jaccard index) was twice as strong as among E- species (mean E+ Jaccard index = 0.032 vs. mean E- Jaccard index = 0.014, a significant difference p = 0.001; Figure S13). These patterns were visualized using richness maps for endophyte-positive and endophyte-negative species (Figure 1b) to highlight differences and similarities in spatial distributions.

*Poa* species with endophytes are more likely to occur in regions with lower precipitation (mean annual precipitation = 668 mm) than those without endophytes (mean annual precipitation = 901 mm), indicating that precipitation strongly influences endophyte distribution. Specifically, we assessed the underlying factors responsible for determining *Epichloë* presence by extracting niche centroids of key climate and soil variables (mean annual temperature, annual precipitation, soil nitrogen, and soil clay content) from each species’ range and tested their relationships with endophyte presence. Bayesian multivariate logistic regressions found that annual precipitation has an influence on endophyte presence (95% Confidence Interval (CI) (-2.23, -0.11); Table S5) and supports the well-established benefit of *Epichloë* endophyte-mediated drought tolerance playing a consistent role in species distributions across this large, ecologically-important grass clade. We also considered the influence of spatial structure of *Epichloë* endophytes on these patterns using a Bayesian spatiophylogenetic model, which confirmed the influence of precipitation on *Epichloë* presence (Figure 1c; Table S6-S7).

### Epichloë endophytes double host species ranges and increase niche breadth

Using phylogenetic generalized least squares regressions, we uncovered that symbiotic *Poa* species have ∼2.5 times larger geographic ranges (mean range size = 6,231,013 km^2^) than those without endophytes (mean range size = 2,533,193 km^2^; PGLS: β = 0.65 ± 0.15, p < 0.0001; model λ = 0.49, n = 105; Figure 2a). In agreement with this finding, the range on the primary niche axis is ∼1.38 times broader for species with endophytes (mean range on primary niche axis = 1.91 SD) than those without (mean range on primary niche axis = 1.39 SD, PGLS: β = 0.15 ± 0.06, p = 0.01; model λ = 0.09, n = 105), suggesting that endophytes play a role in increasing niche breadth in *Poa*. Interestingly, no significant relationship was detected between endophyte presence and environmental hypervolume (Figure 2a; PGLS: β = 0.15 ± 0.12, p = 0.22; model λ = 0.19, n = 105), which could imply that while some niche axes are expanded, endophytes may contract the niche on axes that do not strongly influence species ranges. These results are robust to methodological decisions, remaining consistent when latitude is included as a covariate (Table S8) and when we include a spatial random effect alongside the phylogenetic effect in analyses (Tables S6, S9).

**Figure 2.**
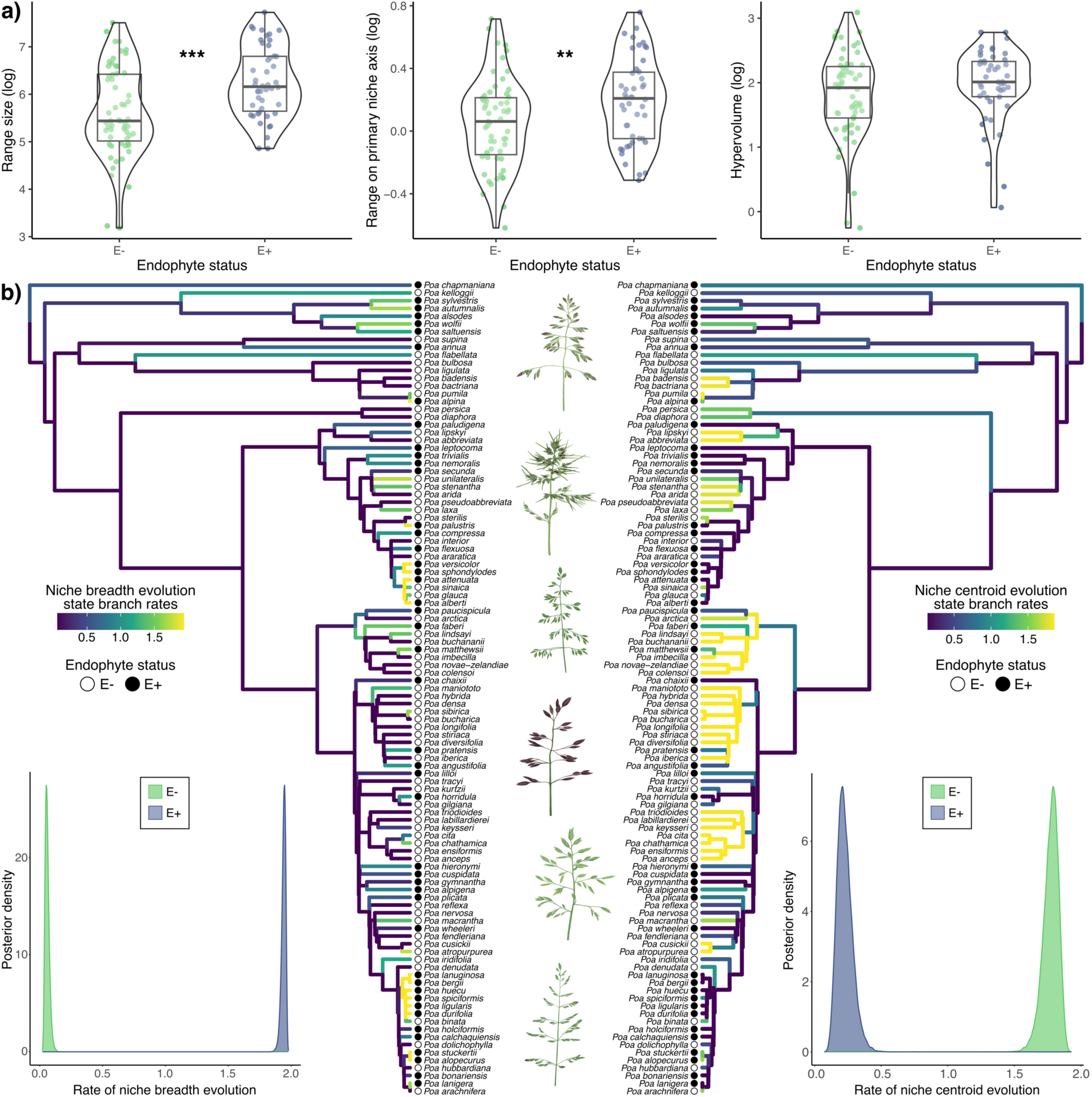
Influences of endophytes on *Poa* niche breadth and niche evolution. A) Results from PGLS models testing the effect of endophytes on geographic range size, range on the primary niche axis, and hypervolume (left to right), stars indicate significance: *** p < 0.0001, ** p = 0.01. B) Displays the *Poa* phylogenetic tree, with state-dependent rates of evolution from MuSSCRat models displayed on the branches for niche breadth (left) and niche centroid (right). Dots at the tips of the tree indicate endophyte status (white = no endophyte, black = endophytes). Density plots next to each tree display estimated rates of evolution. Illustrations by Eleanor Hay.

Interestingly, these mutualists also influenced the evolution of host species’ niches, with mutualistic lineages showing accelerated niche breadth evolution. To assess this, we used Bayesian state-dependent, relaxed, multivariate Brownian motion models of trait evolution (MuSSCRat)^45^ to model how endophyte presence influences the evolution of niche breadth or niche centroid variables, while also considering background rate variation. We constructed two separate models, one testing how endophyte status influences multivariate niche breadth (geographic range size, range on the primary niche axis, and hypervolume) evolution, and the other assessing influences on multivariate niche centroid (mean annual temperature, annual precipitation, soil nitrogen, and soil clay content) evolution. These models show that endophytes have a highly probable influence on rates of niche breadth evolution (posterior probability = 1.0). Rates of niche breadth evolution are over 30 times higher in *Poa* species with endophytes than in species without endophytes (Figure 2b; mean rate of niche breadth evolution_E-_ = 0.06, 95% highest posterior density (HPD) = (0.03, 0.094); mean rate of niche breadth evolution_E+_ = 1.94, 95% HPD = (1.91, 1.97)), indicating that endophyte associations substantially enhance environmental niche breadth lability.

We also found that niche centroid evolution is influenced by endophytes (posterior probability = 1.0). However, in this case, rates of evolution of niche centroids decrease for species that have known associations with *Epichloë* endophytes (Figure 2b; mean rate of niche centroid evolution_E-_ = 1.77, 95% HPD = (1.66, 1.88); mean rate of niche centroid evolution_E+_ = 0.227, 95% HPD = (0.121, 0.345)), implying that endophytes constrain evolution in the mean niche of their hosts. Together, these findings suggest that endophytes promote flexibility within existing niches but may constrain exploration of novel environmental conditions. These patterns and relative rate differences were consistent across different priors on the number of background rate shifts, supporting the robustness of our results (Table S10; Figures S14-S15).

### Fungal endophytes slow diversification in some lineages

Using a Bayesian hidden state speciation and extinction model (HiSSE)^46,47^, we found evidence that endophytes can slow diversification of their host plants (Figure 3). This analysis, which directly tests the influence of endophytes on diversification while controlling for background heterogeneity, supports divergence in diversification rates between E- and E+ species when lineages are in hidden state A (*T*_A_ 95% Credible Interval (CI) = (0.11, 3.77)), but not in hidden state B (Figure 3c; *T*_B_ 95% CI = (-0.19, 0.77)). Additionally, we found that diversification rates differ between hidden states regardless of endophyte status (Figure 3d; T_E-_ 95% CI = (2.92, 4.41); T_E+_ 95% CI = (0.81, 3.23)). These findings suggest that there are distinct modes of diversifications across the *Poa* phylogeny: when lineages are in hidden state A (younger lineages with higher mean diversification rates), symbiotic species are associated with a decrease in diversification, while there is no probable difference in diversification when lineages are in hidden state B (older lineages with lower mean diversification rates). This finding highlights that endophytes influence diversification of some *Poa* lineages but not others. Further, these results support that endophytes constrain speciation, but that this effect is only detectable when diversification rates are high. Alternatively, another character may change between the two hidden states that alters the endophyte’s influence on speciation.

**Figure 3.**
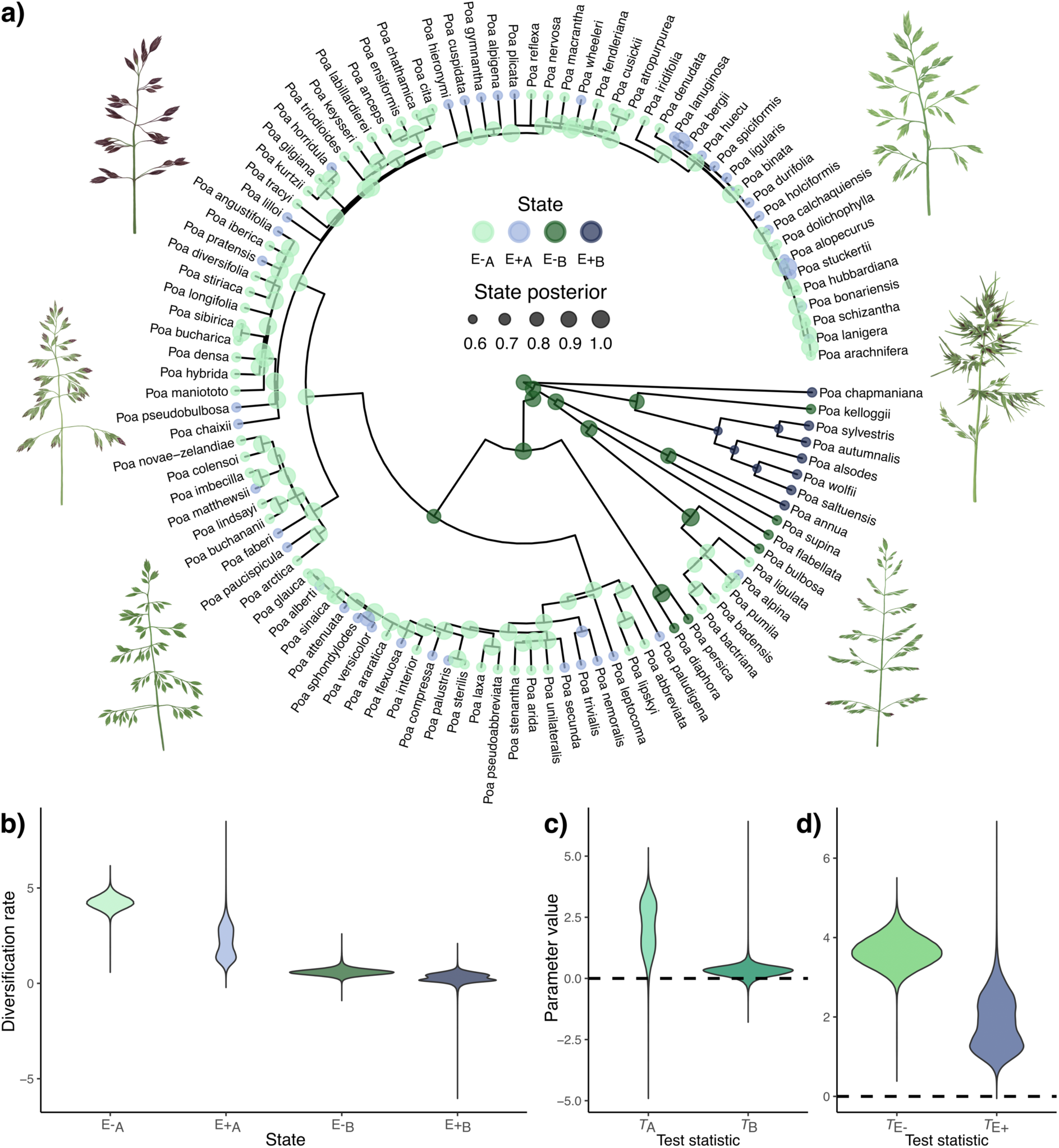
Influences of endophytes on diversification of *Poa*. The phylogeny (a) shows ancestral character history of endophytes from our HiSSE model, with the green colors being E-and blue E+. Panel (b) shows the posterior distribution of net diversification for each of the states. Panel (c) shows the test statistic of the difference in diversification rate between E- and E+ species within each hidden state, with E+ status leading to a decrease in diversification when linages are in hidden state A (younger lineages), but not hidden state B (older lineages). Panel (d) shows the test statistic of difference in diversification rates between hidden states A and B for E-and E+ species, demonstrating the association between hidden states and lineage age. Illustrations by Eleanor Hay.

## DISCUSSION

In this study, we tested the hypothesis that mutualisms increase niche breadth and drive diversification, using the *Epichloë*-grass symbiosis as a model system. Overall, we found that endophyte partnerships are associated with broader and more labile host niche breadths: endophyte-positive species have 2.5-fold larger geographic ranges, 1.4-fold broader tolerances on the primary niche axis, and 30-fold greater rates of niche breadth evolution. Despite this, we also found that endophytes constrain evolution of niche centroids, and we uncovered two distinct modes of diversification for *Poa*, a faster mode in which endophytes decrease diversification rates, and a slower mode in which diversification is driven by other unmeasured factors. Together, our results highlight a nuanced host-symbiont relationship in which mutualists increase niche breadth, but this effect does not translate to increased diversification rates. Our findings challenge the assumption that broader niches always promote diversification and reveal a more complex role for mutualists in shaping macroevolutionary patterns.

Our findings support the hypothesis that mutualisms expand hosts’ niche breadths: symbiotic *Poa* species have ranges ∼2.5 times larger than non-symbiotic species, and rates of niche breadth evolution are >30 times faster. Importantly, these results support empirical findings from species-level grass-*Epichloë* studies. For example, Afkhami et al.^34^ showed that in *Bromus laevipes*, a species from a genus closely related to *Poa*, *Epichloë* endophytes expand their host’s range by 20%. Our results provide the first evidence that this type of niche breadth expansion by endophytes is widespread across species. Within species, there is often variation in endophyte prevalence, which is understood to contribute to expanding niche breadth by enabling symbiotic and non-symbiotic populations to occupy distinct environmental conditions. In *B. laevipes,* E+ and E- populations have distinct niches, with the E+ populations occupying more stressful (drier) environments. Similar patterns have been observed within *Poa*. For example, in *Poa annua*, endophyte prevalence increases with absolute latitude, and is thus thought to have facilitated invasion of Antarctica^38^. It could be the case that all E+ species with increased niche breadth consist of a mix of E+ and E- populations that exhibit niche divergence, however further research on how endophyte prevalence varies across populations within and between species is required to confirm this. We not only highlight that endophyte-positive species tend to have broader niche breadths, but also have increased evolutionary lability in niche breadth. The ability to gain and lose endophyte from a subset of the populations across an E+ species’ range may be what allows for rapid fluctuations in niche breadth. Certainly, the gain or loss of a facultative symbiont can occur much more rapidly than evolution of the genomic architecture underlying stress tolerance.

*Epichloë* endophytes are known to ameliorate abiotic stressors, which may underlie the broader niches in endophyte-associating *Poa*, suggesting that the physiological advantages conferred by endophytes scale up to shape clade-level ecological processes. One of the primary benefits conferred by *Epichloë* is drought tolerance. Our analyses showed that the strongest predictor of endophyte presence across *Poa* was precipitation, and that species with endophytes are found in drier areas. This expands on a pattern identified by Żurek et al.^48^, who explored relationships between bioclimatic variables and *Epichloë* colonization across 26 cool season grass species in Poland, and found that endophyte presence was negatively related to precipitation. *Epichloë* endophytes are known to promote a range of physiological adaptations that can assist with drought tolerance. These include higher root biomass to improve water uptake, improved stomatal regulation to reduce water loss, and increased accumulation of solutes to improve water storage^28,29^. Beyond precipitation, endophytes may also influence tolerance of temperature extremes^48–50^, and we found evidence consistent with this idea. *Poa* species associating with *Epichloë* endophytes have 40% broader tolerances on their primary niche axes (i.e., the niche dimension that is most important for determining host species ranges), which were most often temperature variables (observed in 31 of the 47 endophyte-positive *Poa* species; Table S4), suggesting that *Epichloë* endophytes likely contribute to expanded temperature tolerances. Adaptations to temperature have been identified as important in general for the evolution of grasses^51^ and have been hypothesized to play a key role in the radiation of *Poa*^36^. It is possible that the same physiological mechanisms through which endophytes reduce water loss also improve thermoregulation, enabling hosts to persist under extreme or variable temperature regimes.

Despite the evidence that endophytes expand niche breadth and increase rates of niche breadth evolution, we found that *Epichloë* constrain niche centroid evolution of their hosts. Endophyte-positive *Poa* species exhibit more similar niches per unit of evolutionary time, suggesting that these symbiotic interactions are biased toward certain environmental conditions. *Epichloë* are obligate mutualists: they cannot persist without the host plant. There is growing evidence that obligate relationships tend to stabilize or constrain host species’ niches^52,53^. This could be because the symbiont prefers the ancestral conditions in which the mutualism evolved, and negative fitness consequences of losing the symbiont would constrain evolution into new regions of niche space. At the population-level, *Epichloë* endophytes have varying effects among different life stages of the host^33^, and altering germination and growth success in certain habitats has played a role in modifying niche dimensions in *Poa* species^37^. For example, in *Poa leptocoma* (E+), endophyte presence increased aboveground biomass by 40% under non-flooded conditions and conferred no benefit under flooding, generating directional selection on the mean niche of the host species^54^. In *Bromus laevipes*, range expansion similarly occurred into drier areas, broadening niche breadth along the aridity axis, and increasing range size, but effects were not consistent on all axes, illustrating directional rather than multidimensional niche expansion^34^. Constraints on the niche could explain why we detected no significant influence of endophytes on niche hypervolume. Together, these findings suggest that while endophytes expand some axes and aspects of their hosts’ niche breadths (range size and primary niche axes, which were largely temperature variables), they might constrain other axes.

Although *Epichloë* endophytes expand niche breadth and accelerate rates of niche breadth evolution, we found that these ecological benefits do not translate into increased diversification of *Poa*. Instead, our hidden state speciation and extinction model suggests there are two modes of diversification within the genus: one in which endophytes are associated with reduced diversification rates (primarily in younger lineages), and another mode in which diversification is driven primarily by other, unmeasured factors (primarily in older lineages). Thus, *Poa* provides an example in which mutualists increase ecological niche breadth and lability without promoting lineage diversification. This challenges the assumption that broader niches necessarily lead to higher diversification rates and contrasts with past studies that have typically found that mutualisms promoting niche expansion also accelerate diversification^16,17^. At the same time, analytical models and simulations have demonstrated that obligate mutualisms constrain diversification^22,55^, and a negative effect of mutualists on host diversification has been found in some instances^19^. For example, in deep-sea symbiotic mussels, adaptive radiation was not driven by the evolution of novel symbioses as expected and chemosymbiotic mussels had slowed diversification rates^56^. Similar patterns have been observed in ants that provide defense to plants in exchange for food and shelter: the mutualism evolved more frequently in rapidly diversifying ant lineages, but then subsequently slowed diversification^57^. Taken together, these results suggest that the uncoupling of benefits provided by mutualists and diversification rates could be a common macroevolutionary outcome across the Tree of Life.

The reductions in diversification associated with *Epichloë* may stem from evolutionary constraints imposed by vertically transmitted endophytes and variation in the costs and benefits of hosting symbionts. When considered alongside our niche analyses, these results indicate that endophytes can promote persistence across broader environmental gradients while constraining ecological divergence among lineages. Specifically, while endophytes expand niche breadth and increase rates of niche breadth evolution, they simultaneously constrain niche centroid evolution, limiting hosts to particular regions of environmental space. Such constraints may reduce opportunities for lineage splitting, particularly during periods of rapid diversification. Moreover, all symbiotic interactions incur costs^58^ and empirical studies have shown that fungal seed endophytes have negative effects on their hosts under certain conditions^59–63^. For instance, fungal seed endophytes have been found to have negative impacts on growth, reproduction, and germination success of seeds of *Festuca arizonica,* and the extent of these effects depends on host genotype^59^. Consequently, the direction and magnitude of endophyte effects are understood to be context dependent, varying with the host species, genotype, and environment^64,65^. Such variability in the direction and strength of selection by endophytes on their hosts likely underscores the heterogeneous macroevolutionary dynamics observed across the *Poa* phylogeny.

Understanding how mutualistic interactions shape evolution of their hosts is a central question in ecology and evolution. Overall, our study shows that mutualistic fungal seed endophytes expand ranges, broaden niche breadths, and accelerate niche breadth evolution across *Poa*, but that these benefits do not lead to increased diversification. Instead, depending on lineage, diversification was either unaffected or reduced by *Epichloë* endophytes, highlighting that mutualisms can promote ecological flexibility while constraining macroevolutionary outcomes. Our findings reveal a decoupling of niche expansion from diversification and underscore the complex, context-dependent role of microbial symbionts in shaping evolutionary trajectories.

## METHODS

### Host-endophyte database

We compiled a database of host-endophyte associations using two complementary approaches. 1) a comprehensive literature search to identify which *Poa* species have endophytes based on published records, and 2) a broad screening of *Poa* seeds from germplasm accessions of species not previously tested for the presence of endophytes. When evaluating endophyte status of *Poa* species from the literature, we used the World Checklist of Vascular Plants (WCVP; v13), which recognizes 606 species of *Poa*^66,67^, and searched for any information on endophyte status for each species. We searched both Web of Science and Google Scholar, and for each *Poa* species we used the search strings (“*SPECIES NAME*” OR “*SYNONYM*”) AND (“endophyte” OR “epichlo*” OR “neotyphodium” OR “acremonium”). The Taxonomic Name Resolution Service (v5.3.1)^68,69^ was used to suggest synonyms and match species names from the literature to the WCVP. Because the published literature could be biased toward reporting species with endophyte associations, we contacted authors for unpublished data when broadscale surveys were conducted and not all endophyte-negative plant species were reported. From our literature search, we obtained information on endophyte status for 73 species of *Poa* from over 40 studies, and acquired data on a further 6 species from contacting authors (Table S1 and Table S2).

For our second approach, screening seeds of *Poa* species to detect the presence of seed-borne endophytes, we used the United States Department of Agriculture Germplasm Resources Information Network (grin-global.org) and requested all accessions available for any species of *Poa* lacking endophyte data in the literature. We sampled 15 seeds per species, spread across the available accessions. For species with more than 15 accessions, we prioritized more recent collections (which are more likely to retain endophytes during storage and plant propagation) and geographically distinct locations. A total of 152 seed accessions from 37 *Poa* species were scored for endophyte presence/absence (Table S3). Seeds were softened overnight in a 5% sodium hydroxide solution at room temperature. They were then stained with aniline blue-lactic acid dye and visually assessed under a compound microscope^70^. *Poa* species were scored as endophyte-positive if any assessed seeds contained stained endophyte hyphae within the aleurone layer. This method has been found to yield similar results to immunoblot assay or PCR detection methods^71^, is the method used in most of the published studies included in our literature search, and is a commonly used approach for determining endophyte presence^72^. Of the 37 species screened, 7 were classified as endophyte-positive (Table S3). This positive rate of 22% falls in line with estimates that fungal seed endophytes occur in 20 – 30% of grasses^23^.

### Environmental and niche breadth data

We used three metrics to assess niche breadth across *Poa* species: geographic range size, range on primary niche axis, and niche hypervolume. Geographic range size and range on the primary niche axis were estimated using species distribution modeling, and hypervolumes were calculated in multivariate environmental space following Blonder et al.^43^.

We downloaded occurrence records from the Global Biodiversity Information Facility (GBIF) for each *Poa* species we had endophyte status for using the *rgbif* package^73^ in R^74^. Occurrence records were filtered using the *CoordinateCleaner* package^75^, which flags temporal and spatial errors commonly found in the GBIF database. To do this, we used the function ‘*clean_coordinates’* and included all coordinate cleaner tests except for the “country” and “outliers” tests, because preliminary checks indicated that these tests were removing accurate localities. All remaining occurrence records were then manually plotted and checked against known ranges following Plants of the World Online^76^, and any further erroneous records that fell far outside species’ known ranges were removed manually. We retained only species with at least 15 unique occurrence points to ensure robust niche breadth estimation^77^: from our 116 species with endophyte data, we found that 112 *Poa* species had sufficient occurrence data after cleaning (mean = 8629, range = 15 – 228,684; Table S4).

Raster layers for 19 climate variables at a resolution of 2.5 arc minutes were obtained from WorldClim^78^. Soil layers (nitrogen, pH, soil organic carbon, clay content, and sand) were obtained from www.soilgrids.org^79^ at 1 km resolution. Soilgrids layers are measured at different depths (0 – 5 cm, 5 – 15 cm, 15 – 30 cm, 30 – 60 cm, 60 – 100 cm, and 100 – 200 cm). Clay content, pH, and sand are highly correlated across the different depths (Pearson correlation coefficients > 0.84), therefore we selected the top layer (0 – 5 cm) to represent these variables. For nitrogen and soil organic carbon content, we used the mean of the top three layers (0 – 5 cm, 5 – 15 cm, 15 – 30 cm) because past studies have found the majority of root density of many grass species, including *Poa* species, to be within the top 30 cm of soil^80^. Soil layers were aggregated to the same grid as the climate data using the ‘*aggregate’* function from the *terra* R package^81^. All raster layers were then standardized to have a mean of 0 and standard deviation of 1 to ensure niche breadth on each environmental layer was comparable.

To assess geographic range size and range on the primary niche axis, we constructed niche models using MaxEnt v3.4.3 in R using the ‘*MaxEnt’* function from the *predicts* package^42^. We used the cleaned *Poa* occurrence records and all raster layers as input data. To determine the geographic extent for each model to be calibrated on, we generated species-specific buffer distances around each *Poa* species’ point occurrence records. We followed the approach used by Mothes et al.^82^ and determined appropriate buffer distances by calculating the Euclidean distance between the two most spatially segregated clusters of localities for each species. The resulting buffer includes the area the species would have historically occurred across, or dispersed across over evolutionary time, and assumes the species would currently be present there if the conditions were suitable.

MaxEnt is a presence-only algorithm and by default chooses random background points to act as pseudo-absences from within the geographic extent. This approach is problematic because background points could randomly be chosen from areas of suitable habitat that are generally inaccessible to researchers, and where the presence of the species has thus not been determined. To account for this sampling bias, we used target-group background points^83^. This approach uses localities of similar species to better reflect true absences, assuming if researchers in the field identified similar species at that locality, they likely would have identified the study species if it were present; this method has been proven to improve MaxEnt model estimates^84^. We used all grasses (Poaceae) as our target group and downloaded occurrence records from GBIF and filtered using the *CoordinateCleaner* package^75^. We then thinned the occurrence records to one record per 2.5 minute grid cell using the ‘*elimCellDuplicates’* function from the *enmSdmX* R package^85^.

To prevent potential overfitting of the MaxEnt models, we performed model selection to choose the best beta multiplier for MaxEnt’s built-in regularization procedure, which balances model fit and model complexity. To identify the optimal regularization multiplier, we followed Warren and Seifert^86^ and generated a total of 24 models per species with regularization multipliers ranging from 0.2 to 1 in increments of 0.2 and integers 1 – 20. We chose the best model based on AIC_c_ and ran the final model using the selected regularization multiplier, 10-fold cross validation (training on 90% of occurrences and testing on the remaining 10% per fold), and the default feature settings. Model performance was assessed using various metrics. We assessed overall fit using the area under the Receiver Operating Characteristic curve (AUC). This represents the ability of the model to differentiate between suitable and unsuitable habitat. We also calculated AUC_DIFF_, which is the difference in the AUC values between the test and training data averaged across folds, and provides a measure of model overfitting (Table S4)^86^.

From the final MaxEnt model for each *Poa* species, geographic range size and range on the primary niche axis were extracted (Table S4). Geographic range size was estimated based on suitability maps using the maximum test sensitivity and specificity (maxSSS) threshold^87^. To estimate range on the primary niche axis, we identified the environmental variable with the greatest permutation importance, since this metric has been shown to accurately identify environmental factors underlying species persistence^88^, and the range on this axis was calculated by extracting minimum and maximum values across the geographic range. Because all raster layers were standardized prior to analysis, the ranges on all environmental variables are comparable. We also generated richness maps for endophyte-positive and endophyte-negative species from the range maps to visualize spatial patterns of endophyte presence across *Poa*. To assess differences between richness maps, we quantified geographic range overlap among *Poa* species to test whether endophyte-positive (E+) species exhibit greater overlap than endophyte-negative (E-) species. Pairwise species overlap was calculated using the Jaccard index, and the mean overlap was calculated separately for E+ and E- species. Significance was assessed using 5,000 permutations in which endophyte labels were randomly shuffled across species, preserving group sizes. One-sided p-values were computed as the proportion of permuted test statistics greater than or equal to the observed test statistic, with the test statistic being the mean Jaccard index for E+ species minus the mean Jaccard index for E- species.

To calculate hypervolume, we used the same standardized climatic and soil variables and extracted their values at all the known *Poa* occurrence points. Principal component analysis was used to reduce the dimensionality of this data using the ‘*prcomp*’ R function. We retained the first 4 PCs, which explained 84.3% of the total variation (Table S11) and used these four environmental axes to determine hypervolumes for each species. Hypervolumes were estimated for each species using the *‘hypervolume’* function from the *hypervolume* package^89^ with a Gaussian kernel density estimation.

To understand what environmental conditions are associated with *Epichloë* fungal seed endophytes, we also calculated the niche centroid for each *Poa* species. This was accomplished by calculating the mean value across all grid cells within a species’ range (again based on the maxSSS threshold) on the following niche axes: mean annual temperature, annual precipitation, soil nitrogen, and clay content. These niche axes were selected based on being relatively independent (all absolute pairwise correlation coefficients < 0.7) and each representing an important component of climatic or soil niche space. To account for latitudinal variation in range size (Rapoport’s rule)^90^, we similarly estimated latitudinal centroids for each species.

### Phylogeny

We used a range of phylogenetic comparative methods to assess the influence of fungal seed endophytes on *Poa* niches, niche breadth, and diversification. Analyses were conducted using a phylogeny from Elliott et al.^39^. This phylogeny was constructed for the order Poales and contains 45% of grasses (Poaceae) and 280 species of *Poa* (46.2% complete sampling for *Poa*). This tree was constructed using a phylogenomic backbone of 353 nuclear loci (Angiosperms353)^91,92^ combined with a supertree approach to generate trees for groups of families identified within the backbone. The topology is consistent with past studies, and divergence times were estimated with primary and secondary calibrations using a penalized likelihood approach^39^. The estimated crown age of *Poa* in this tree is 5.1 million years. This is slightly younger than some estimates from previous studies^36,93,94^, but falls within the known range for this clade^40,41^. This phylogeny contains 104 of the 116 species we have endophyte status for, and we were able to match 3 additional species to their closest relative in the tree, bringing this total up to 107 species (Table S1).

### Phylogenetic comparative methods

We started by assessing phylogenetic patterns of endophyte presence in *Poa*. We calculated phylogenetic signal of endophytes across *Poa* using the D statistic, a measure of phylogenetic signal for binary traits^95^. This was estimated in R using the ‘*phylo.d’* function from the *caper* package^96^.

To identify environmental factors influencing endophyte presence across *Poa* species, we employed multivariate phylogenetic logistic regression. Models were implemented in R with the package *brms*^97^ for Bayesian models using the probabilistic programming language STAN^98,99^. This approach uses a Markov chain Monte Carlo (MCMC) sampler that uses techniques based on Hamiltonian Monte Carlo^100^. Endophyte presence/absence was modeled using a binomial family and we used niche centroids as predictors. For our models, we pruned the *Poa* phylogeny to the species with both phylogenetic and niche data available (n = 105), using the ‘*drop.tip’* function from the R package *ape*^101^. The ‘*vcv’* function from the *ape* package was used to generate a phylogenetic variance-covariance matrix from the phylogeny to include as a random factor in the model. Models were run for 2000 MCMC iterations with 4 chains and convergence was assessed by ensuring the estimated potential scale reduction factor (Rhat) statistic reached 1.0^99^. Effects were judged as probable if the 95% credible interval of the effect size did not overlap zero.

To account for potential geographical clustering of *Epichloë* endophytes, we decided to fit a spatiophylogenetic model to consider the influence of spatial non-independence alongside phylogenetic non-independence on these patterns. Models were fit using integrated nested Laplace approximation (INLA) in the R package *INLA*^102,103^, which provides accurate and computationally efficient approximations to posterior distributions for latent Gaussian models^103,104^. We followed a modelling framework described by Dinnage et al.^105^ that uses INLA to simultaneously deal with phylogenetic and spatial non-independence, and incorporates fixed and random effect predictors that vary spatially with a response variable measured at the species level. This approach retains variation in the size and shape of species ranges and the extent of overlap between different species^105^. We used this method to model how environmental variables influence endophyte presence and absence, using a “binomial” family with a logit link function. For the phylogenetic effect, INLA requires a phylogenetic precision matrix, which is the inverse of a phylogenetic covariance matrix. Following Dinnage et al.^105^, the phylogenetic covariance matrix was standardized by dividing by its determinant raised to the power of 1/N_species_ prior to inverting. To model the spatial effect, we used a spatial mesh, which was constructed over *Poa* species occurrence records and averaged across the spatial random fields of the occurrence records for each species. This approach integrates a spatial random field across each species’ distribution, so each species contributes a single datapoint to the likelihood, reducing any bias that could arise when the number of occurrence records varies between species (see Dinnage et al.^105^ for further model explanation). We assessed a range of spatial meshes that varied in coarseness (2528, 4688, and 13018 vertices; Figure S16) and final models used a mesh consisting of 2528 vertices. Models were repeated over several phylogenetic and spatial priors to test the robustness of results. Model comparison based on the marginal log-likelihood, deviance information criterion (DIC), and the widely applicable Bayesian information criterion (WAIC) was used to assess the fit of different meshes and priors (Table S7). The probable influence of effects in the models were based on whether the 95% credible interval of the effect size overlapped with zero.

We then used phylogenetic generalized least squares regression (PGLS) to test whether *Epichloë* endophyte presence influences niche breadth. PGLS models were implemented in R using the packages *ape*^101^ and *nlme*^106^, with Pagel’s lambda estimated by maximum likelihood to account for phylogenetic signal in model residuals^107,108^. We performed bivariate models to test for direct associations between endophyte presence and each response variable, followed by multivariate models to also assess the influence of latitude. Niche breadth variables were log-transformed prior to analysis to improve normality and absolute values of latitude were used. We also accounted for the influence of spatial covariance on these patterns and repeated models using INLA to include a spatial random effect alongside the phylogenetic random effect in analysis. We modeled how endophytes influence niche breadth variables with a gaussian family. We used the same approach as described above, with the same spatial mesh, and tested a range of phylogenetic and spatial priors to assess the robustness of results (Table S9).

To assess whether *Epichloë* endophytes influence niche breadth evolution or niche centroid evolution in *Poa*, we used state-dependent, relaxed, multivariate Brownian motion models of trait evolution^45^ (MuSSCRat) implemented in RevBayes v1.2.4^47^. We employed two separate models in which we tested the influence of endophytes on niche breadth evolution and niche centroid evolution. In both models, endophyte status was modeled as a discrete character (presence vs. absence), while in the first model niche breadth traits (geographic range size, range on the primary niche axis, and hypervolume) were used as the multivariate continuous response variables, and in the second model niche centroids (mean annual temperature, annual precipitation, soil nitrogen, and soil clay) were used. The MuSSCRat model is advantageous because it jointly estimates the evolutionary history of the discrete and continuous characters, while also considering background rate variation (i.e., variation in trait evolution unrelated to endophyte status). By allowing evolutionary rates to vary along branches and among continuous characters, MuSSCRat reduces biases in rate estimates. This model also incorporates hypothesis testing by comparing state-dependent and state-independent scenarios using a reversible-jump MCMC algorithm^45^. The MCMC was run for at least 500,000 generations with a 10% burn-in, and convergence was assessed using Tracer v1.7.2^109^ by ensuring effective sample sizes (ESS) were above 200. To evaluate the sensitivity of posterior parameter estimates, we repeated the MCMC across different priors on the number of background rate shifts of the continuous characters (5, 10, and 20 shifts; Table S10). Model results were visualized using the R packages *RevGadgets*^110^ and *ggtree*^111^.

To determine whether *Epichloë* endophytes influence diversification in *Poa*, we used hidden state speciation and extinction models (HiSSE)^46^. HiSSE models extend traditional binary state diversification models by incorporating hidden (unmeasured) states, enabling estimation of speciation, extinction, and net diversification rates while controlling for background heterogeneity^46^. This framework addresses known limitations of earlier state-dependent approaches^112,113^. We performed a Bayesian implementation of HiSSE using RevBayes^47^. Our model included two observed states, absence of endophytes (E-) and presence of endophytes (E+), and two hidden states (A or B), yielding four diversification regimes (E-_A_, E+_A_, E-_B_, E+_B_). We allowed asymmetric transition rates between all observed and hidden states. In this model, if both of the E+ rates are higher than the E- rates, then the analysis would suggest that *Poa* species with endophytes diversify faster than those without endophytes. We compare net diversification rates across these regimes to evaluate whether differences were attributable to the observed trait (*Epichloë*) or to hidden background factors, which represent other unmeasured factors or heterogeneity, but not the character of interest. One benefit to implementing HiSSE in RevBayes is that it can incorporate missing character state data. Because incomplete sampling is known to bias speciation and extinction estimates, we used the *Poa* phylogeny containing 280 species alongside our character state data for 107 species, with all missing species coded as ‘?’ (Figure S17). We then accounted for incomplete taxon sampling by specifying a global sampling fraction of 42.6%, based on an estimated total of 606 *Poa* species from the WCVP. We assumed the state value at the root was uncertain and used a Dirichlet prior with equal frequencies for all state values, which were estimated through the MCMC. Posterior estimates of root state frequencies indicated that the root was most likely in the E-_B_ state (posterior probability = 0.37), with lower support for E-_A_ (0.20), E+_A_ (0.20), and E+_B_ (0.23). We ran two MCMC chains of 100,000 generations with a 10% burnin, and checked for convergence between the chains and ensured all ESS were above 200 using Tracer v1.7.2^109^. We summarized posteriors and plotted results in R using the *ggplot2* package^114^. We followed approaches of previous studies^115,116^ and used our HiSSE results to create test statistics to determine the probability of endophytes influencing the diversification of *Poa*.

## AUTHOR CONTRIBUTIONS

E.M.H. contributed to study conception, conducted all analyses, and wrote the manuscript. R.Z.- F. contributed to HiSSE analyses and code and reviewed the manuscript. M.E.A. contributed to study conception, statistical input, and manuscript feedback. C.A.S. contributed to study conception, species distribution modelling code, statistical input, and manuscript feedback. All authors approved the final manuscript.

## ACKNOWLEGEMENTS

We would like to thank Christopher Schardl, Jenn Rudgers, and Marjorie Weber for helpful discussions on aspects of this work. Additional endophyte survey data was shared by Jenn Rudgers and Leopoldo Iannone. Members of the Afkhami Lab and Conservation Ecology Lab provided helpful feedback on an earlier version of this manuscript. We thank the USDA Agricultural Research Service for providing germplasm through the Germplasm Resources Information Network (GRIN-Global). This research was supported by a National Science Foundation DEB-2030060 awarded to MEA and CAS.

