## Supplementary material for "Mutualistic microbes drive niche expansion yet constrain diversification": Figures S1-S17

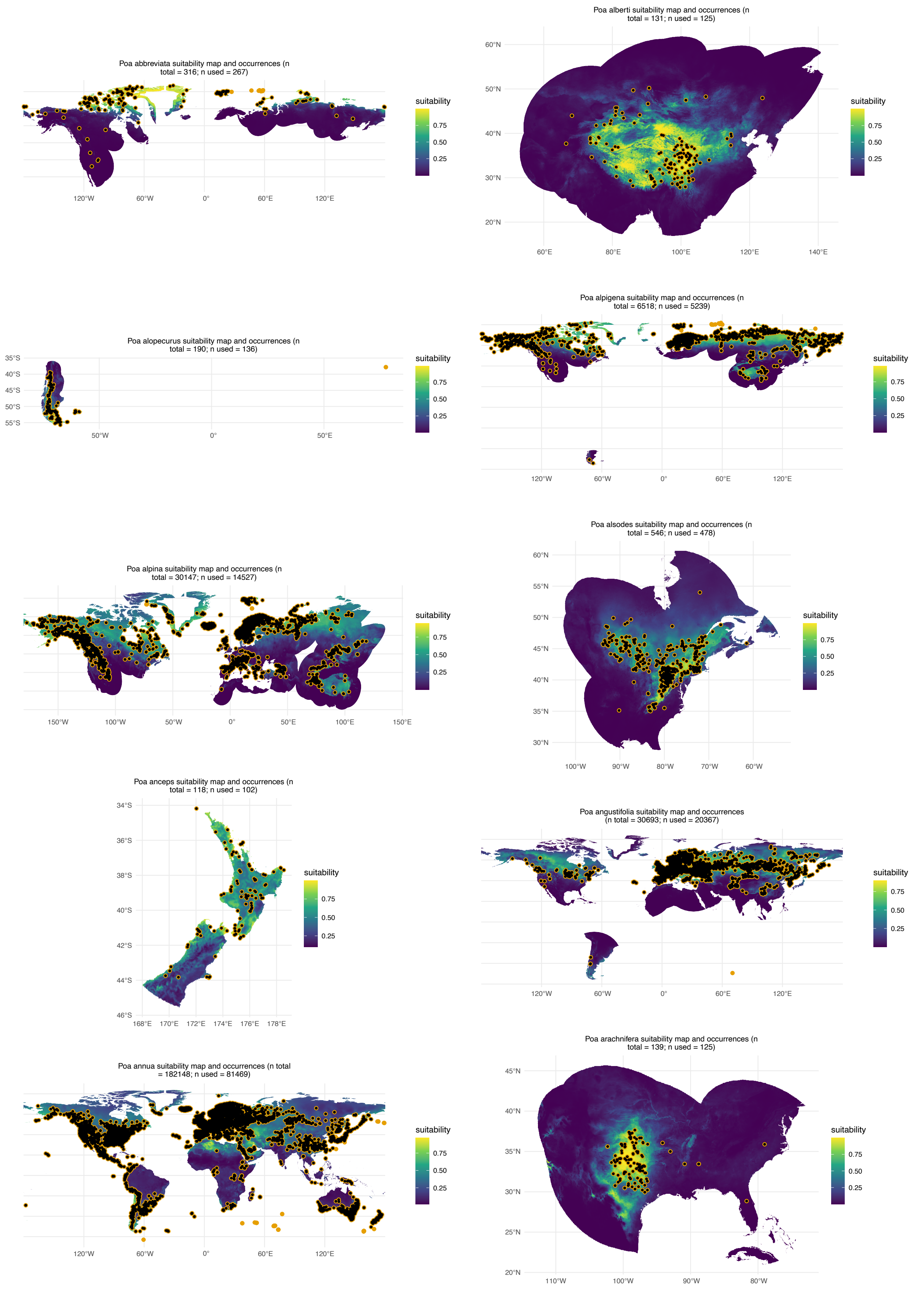


**Figure S1. *Poa* species occurrence records and suitability maps from MaxEnt models.** Geographic range size was estimated based on these suitability maps, using the maximum test sensitivity and specificity (maxSSS) threshold (provided in Table S4). Dots are occurrence records; orange dots represent occurrence records input to the model and dots filled with black indicate occurrence records used by the MaxEnt model.

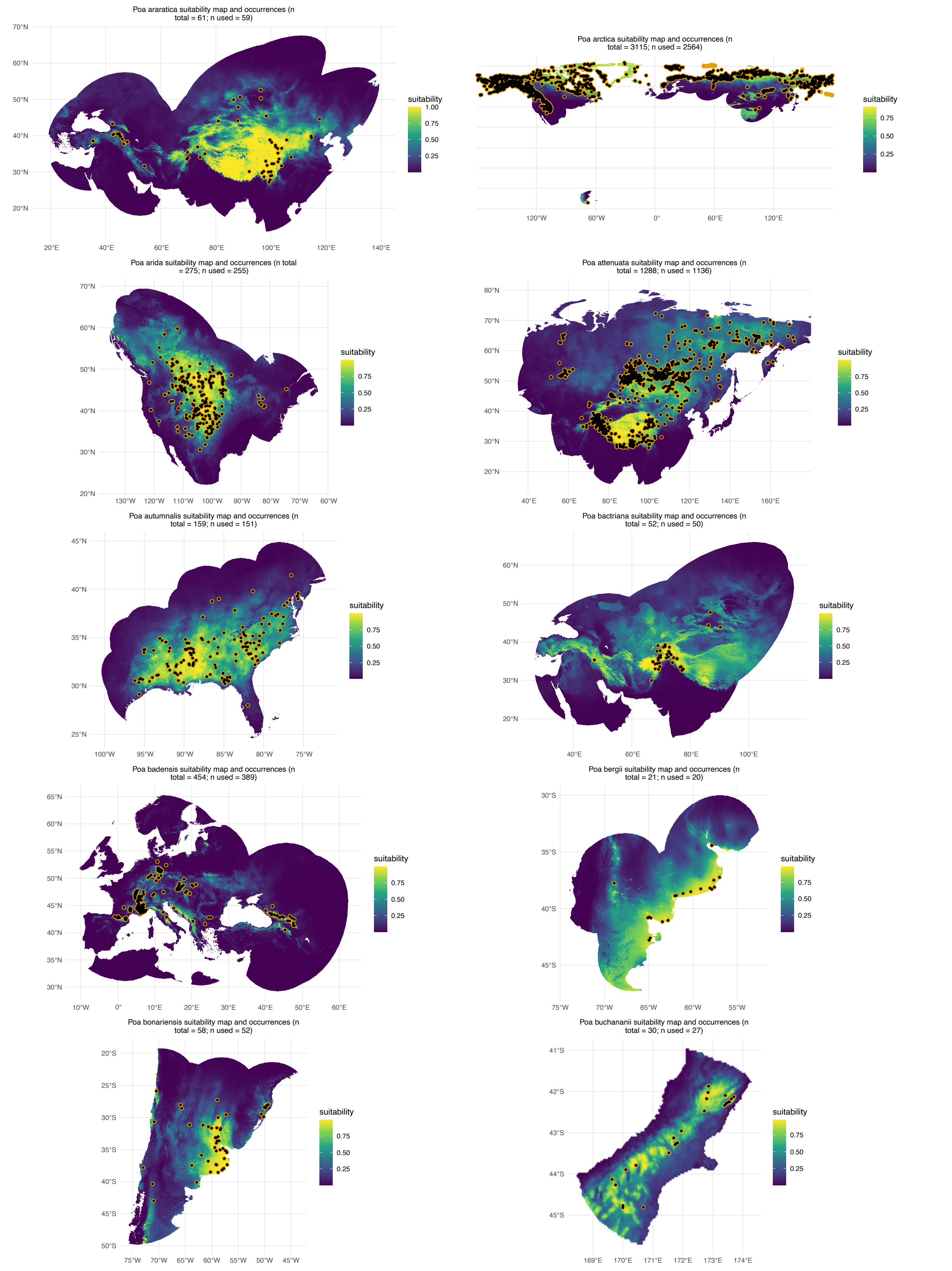


**Figure S2. *Poa* species occurrence records and suitability maps from MaxEnt models.** Geographic range size was estimated based on these suitability maps, using the maximum test sensitivity and specificity (maxSSS) threshold (provided in Table S4). Dots are occurrence records; orange dots represent occurrence records input to the model and dots filled with black indicate occurrence records used by the MaxEnt model.

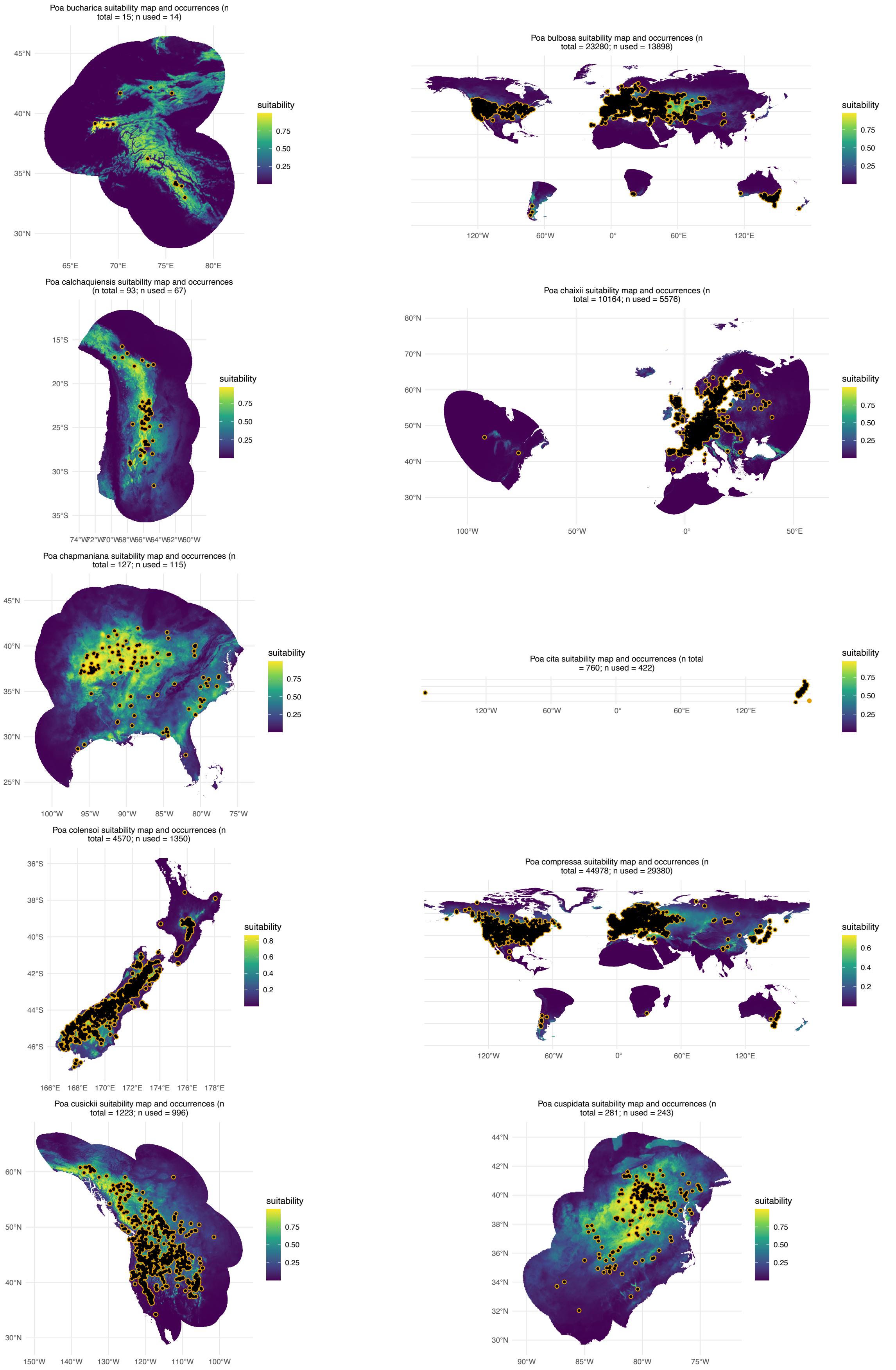


**Figure S3.** ***Poa*** **species occurrence records and suitability maps from MaxEnt models.** Geographic range size was estimated based on these suitability maps, using the maximum test sensitivity and specificity (maxSSS) threshold (provided in Table S4). Dots are occurrence records; orange dots represent occurrence records input to the model and dots filled with black indicate occurrence records used by the MaxEnt model.

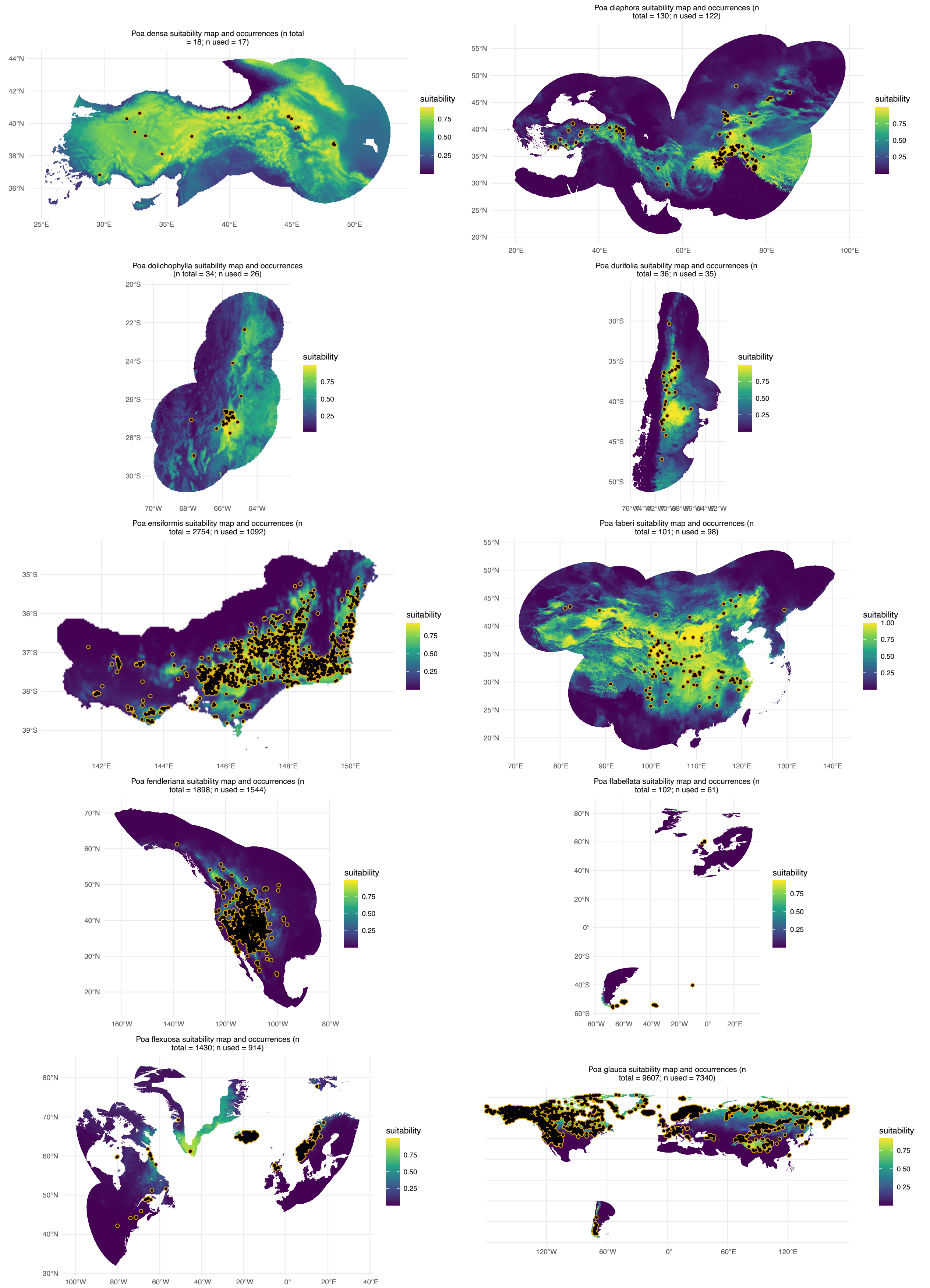


**Figure S4. *Poa* species occurrence records and suitability maps from MaxEnt models.** Geographic range size was estimated based on these suitability maps, using the maximum test sensitivity and specificity (maxSSS) threshold (provided in Table S4). Dots are occurrence records; orange dots represent occurrence records input to the model and dots filled with black indicate occurrence records used by the MaxEnt model.

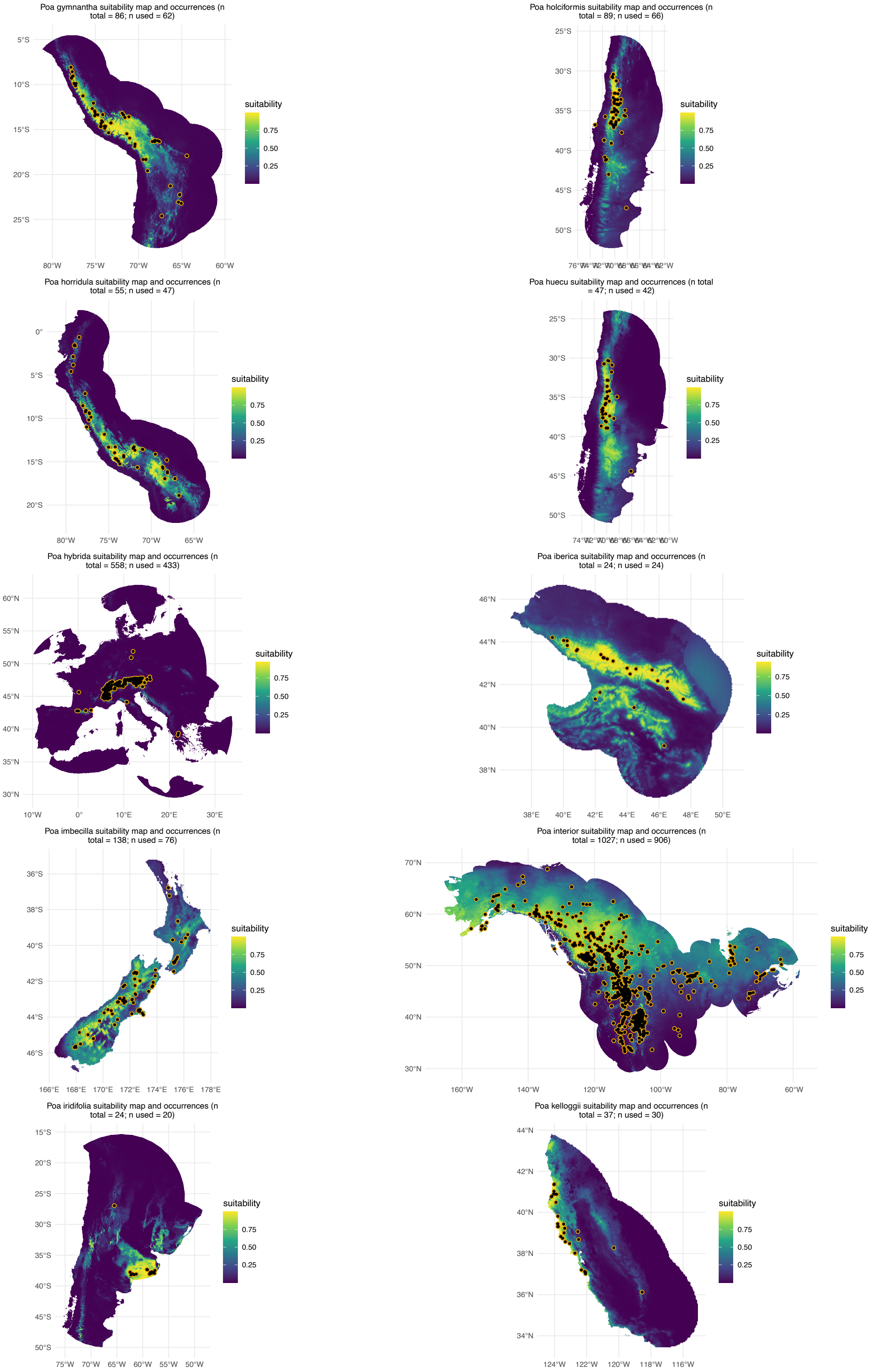


**Figure S5. *Poa* species occurrence records and suitability maps from MaxEnt models.** Geographic range size was estimated based on these suitability maps, using the maximum test sensitivity and specificity (maxSSS) threshold (provided in Table S4). Dots are occurrence records; orange dots represent occurrence records input to the model and dots filled with black indicate occurrence records used by the MaxEnt model.

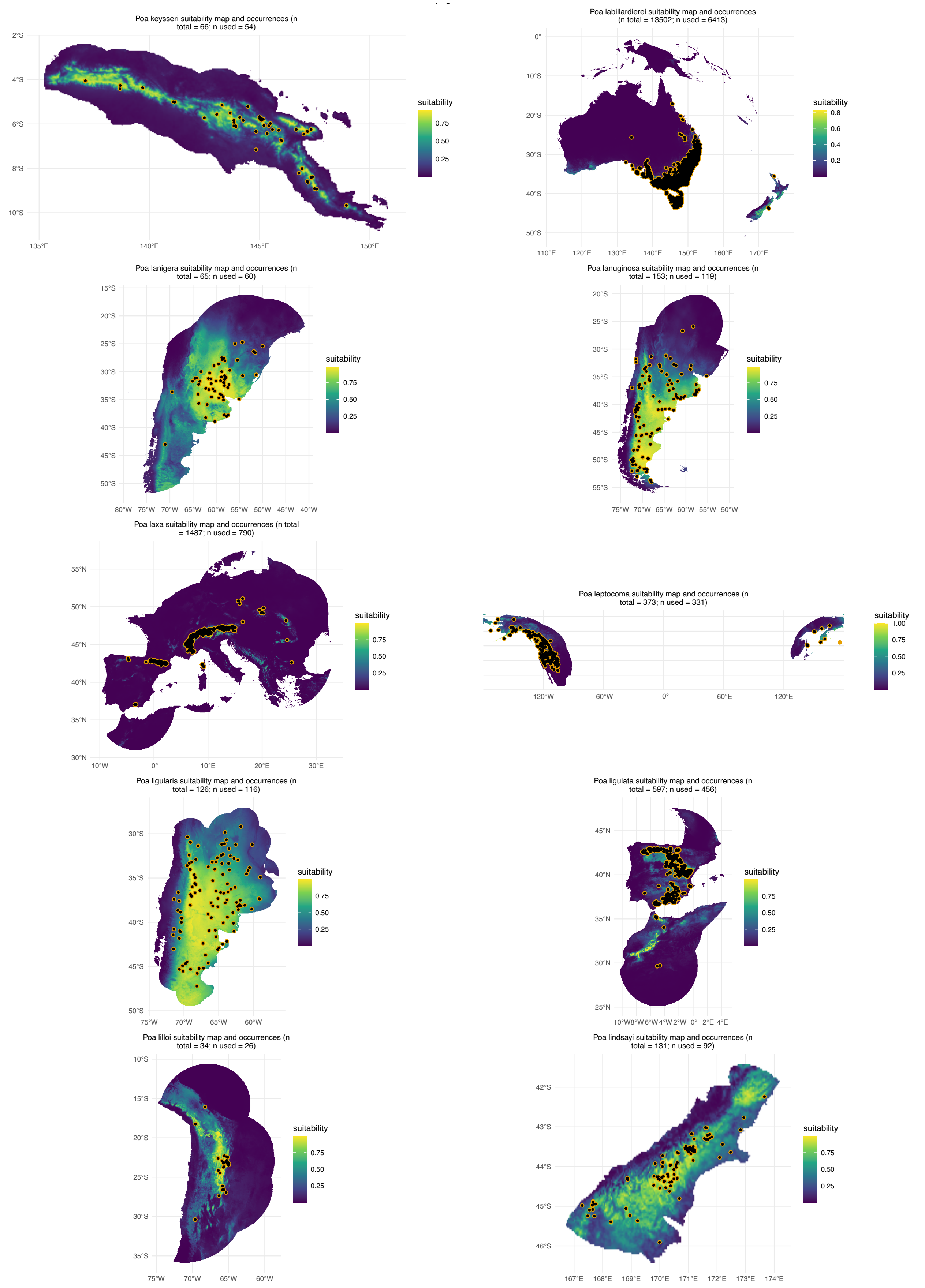


**Figure S6. *Poa* species occurrence records and suitability maps from MaxEnt models.** Geographic range size was estimated based on these suitability maps, using the maximum test sensitivity and specificity (maxSSS) threshold (provided in Table S4). Dots are occurrence records; orange dots represent occurrence records input to the model and dots filled with black indicate occurrence records used by the MaxEnt model.

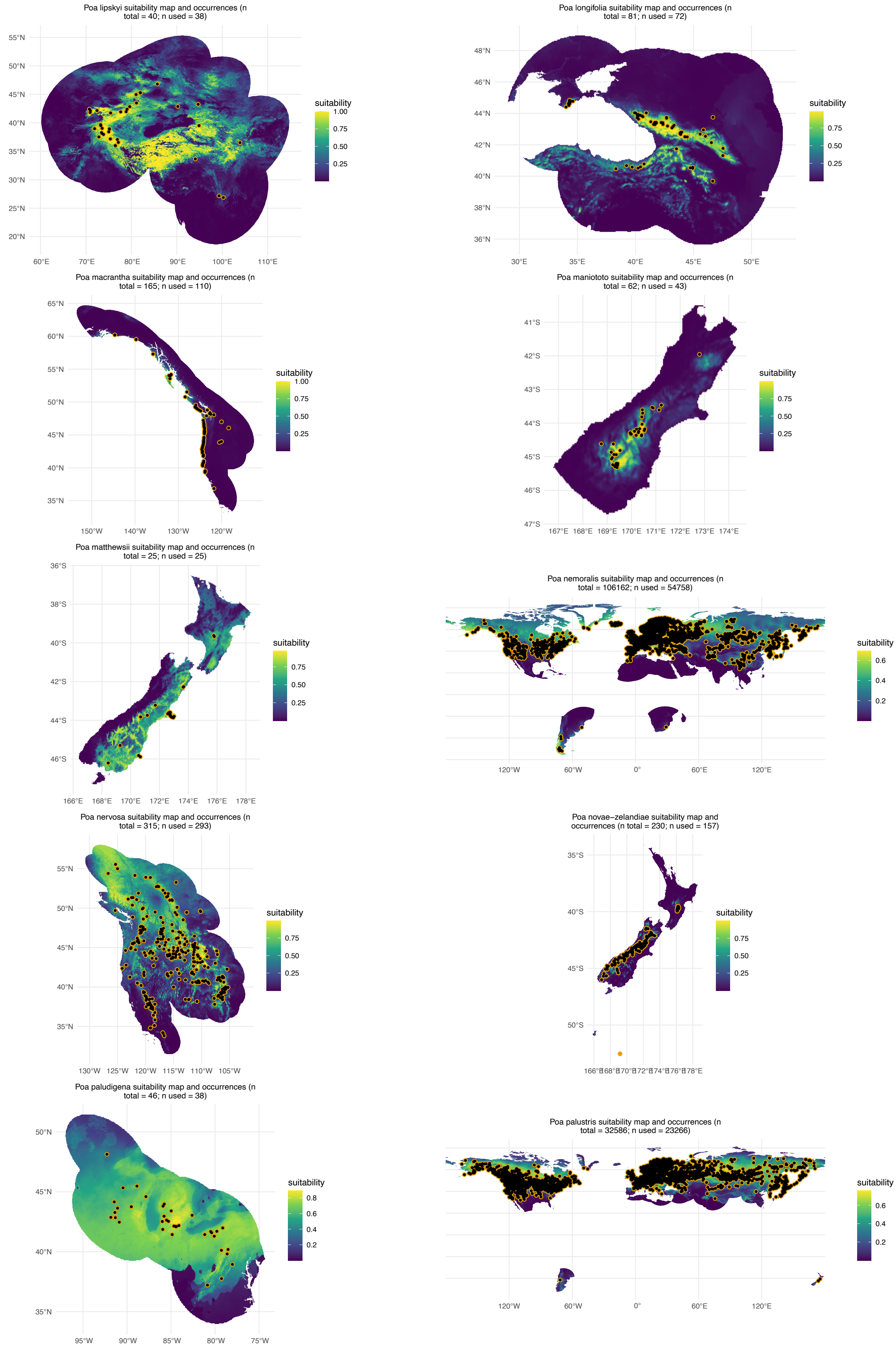


**Figure S7. *Poa* species occurrence records and suitability maps from MaxEnt models.** Geographic range size was estimated based on these suitability maps, using the maximum test sensitivity and specificity (maxSSS) threshold (provided in Table S4). Dots are occurrence records; orange dots represent occurrence records input to the model and dots filled with black indicate occurrence records used by the MaxEnt model.

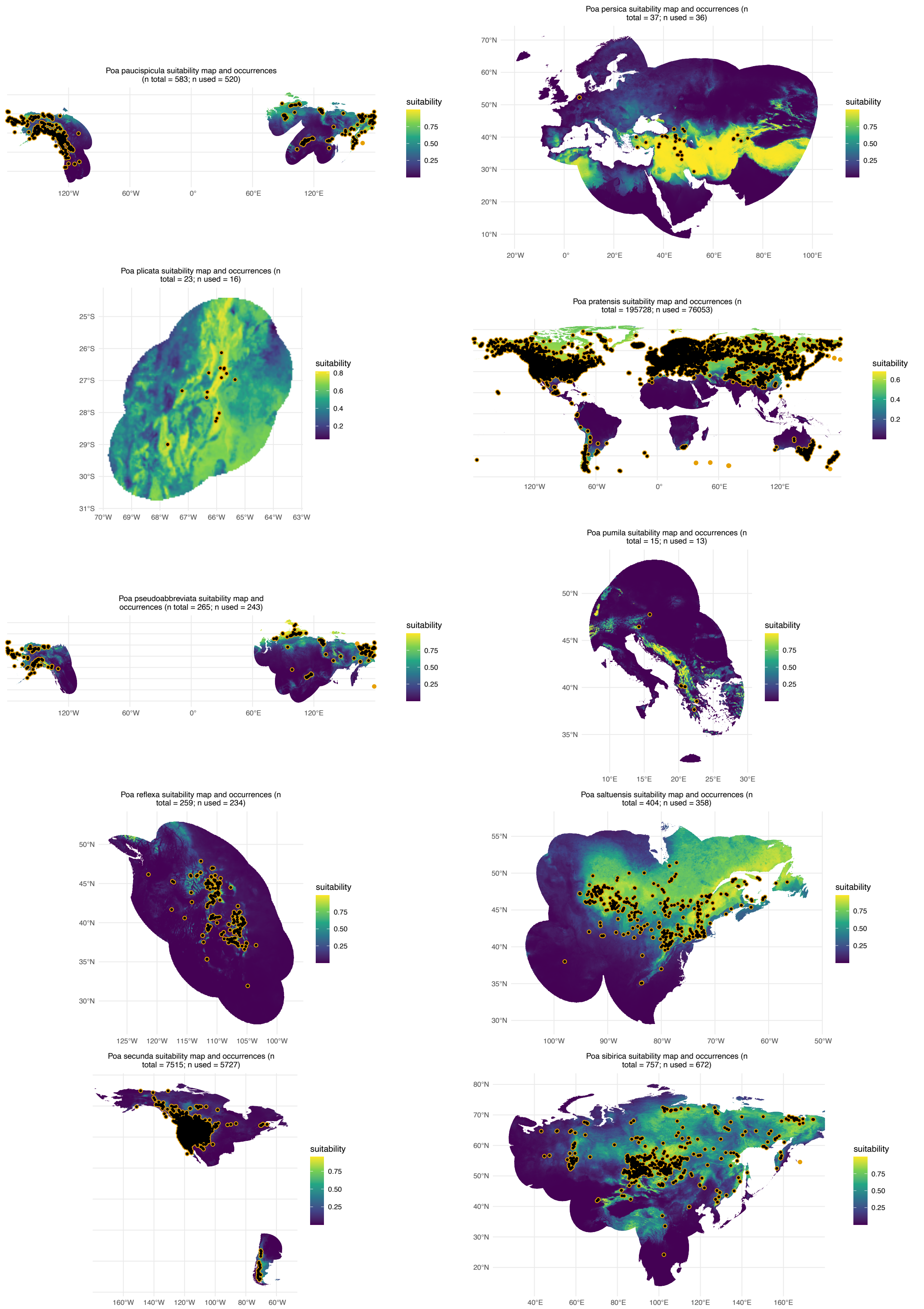


**Figure S8. *Poa* species occurrence records and suitability maps from MaxEnt models.** Geographic range size was estimated based on these suitability maps, using the maximum test sensitivity and specificity (maxSSS) threshold (provided in Table S4). Dots are occurrence records; orange dots represent occurrence records input to the model and dots filled with black indicate occurrence records used by the MaxEnt model.

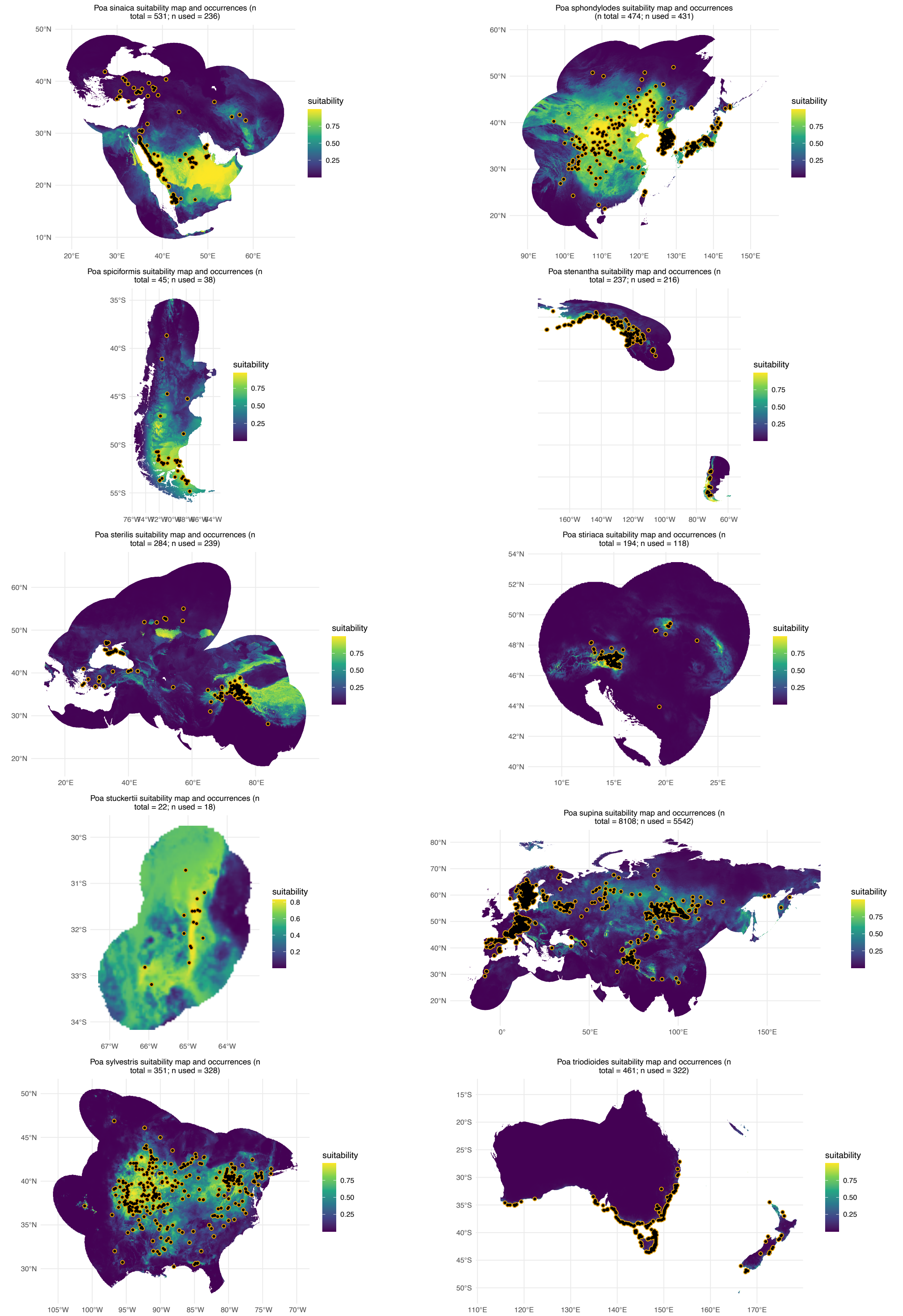


**Figure S9. *Poa* species occurrence records and suitability maps from MaxEnt models.** Geographic range size was estimated based on these suitability maps, using the maximum test sensitivity and specificity (maxSSS) threshold (provided in Table S4). Dots are occurrence records; orange dots represent occurrence records input to the model and dots filled with black indicate occurrence records used by the MaxEnt model.

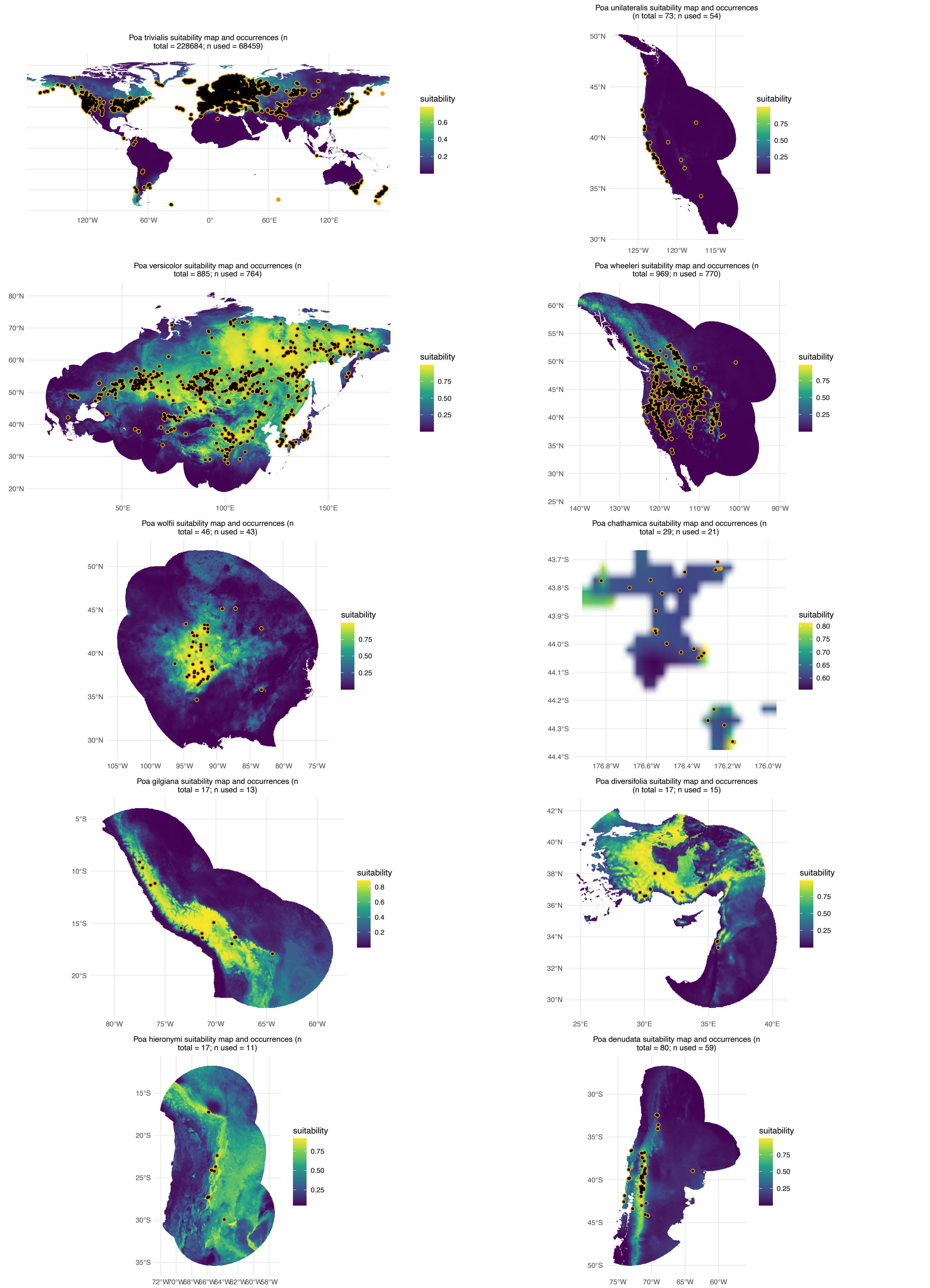


**Figure S10. *Poa* species occurrence records and suitability maps from MaxEnt models.** Geographic range size was estimated based on these suitability maps, using the maximum test sensitivity and specificity (maxSSS) threshold (provided in Table S4). Dots are occurrence records; orange dots represent occurrence records input to the model and dots filled with black indicate occurrence records used by the MaxEnt model.

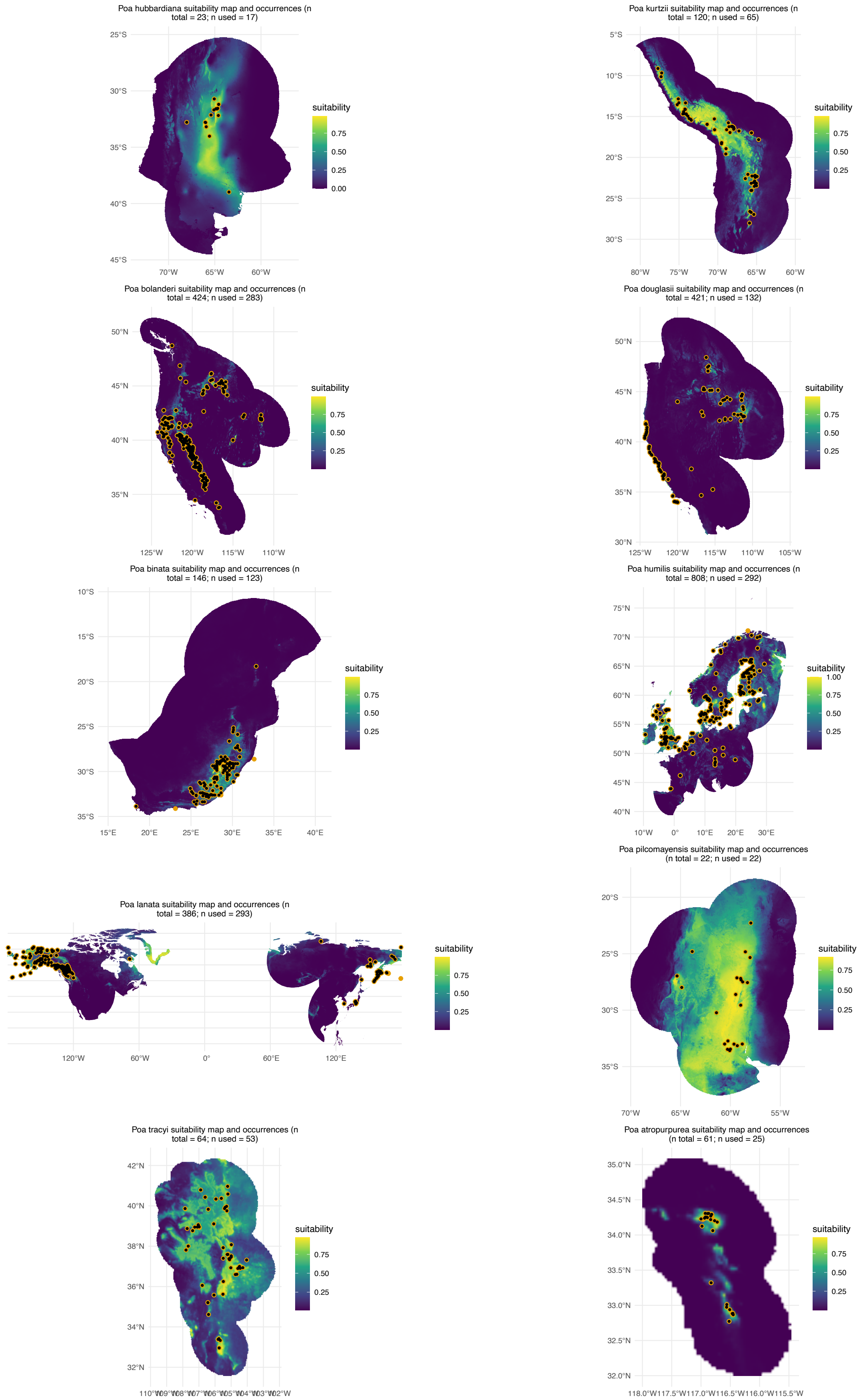


**Figure S11. *Poa* species occurrence records and suitability maps from MaxEnt models.** Geographic range size was estimated based on these suitability maps, using the maximum test sensitivity and specificity (maxSSS) threshold (provided in Table S4). Dots are occurrence records; orange dots represent occurrence records input to the model and dots filled with black indicate occurrence records used by the MaxEnt model.

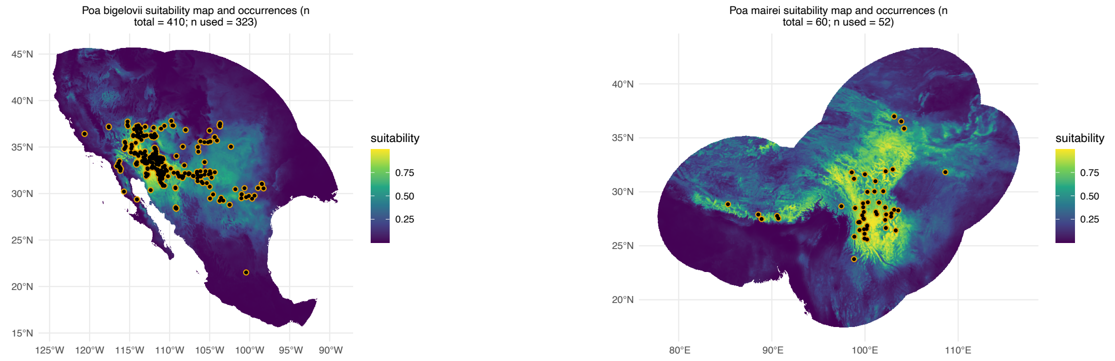


**Figure S12. *Poa* species occurrence records and suitability maps from MaxEnt models.** Geographic range size was estimated based on these suitability maps, using the maximum test sensitivity and specificity (maxSSS) threshold (provided in Table S4). Dots are occurrence records; orange dots represent occurrence records input to the model and dots filled with black indicate occurrence records used by the MaxEnt model.

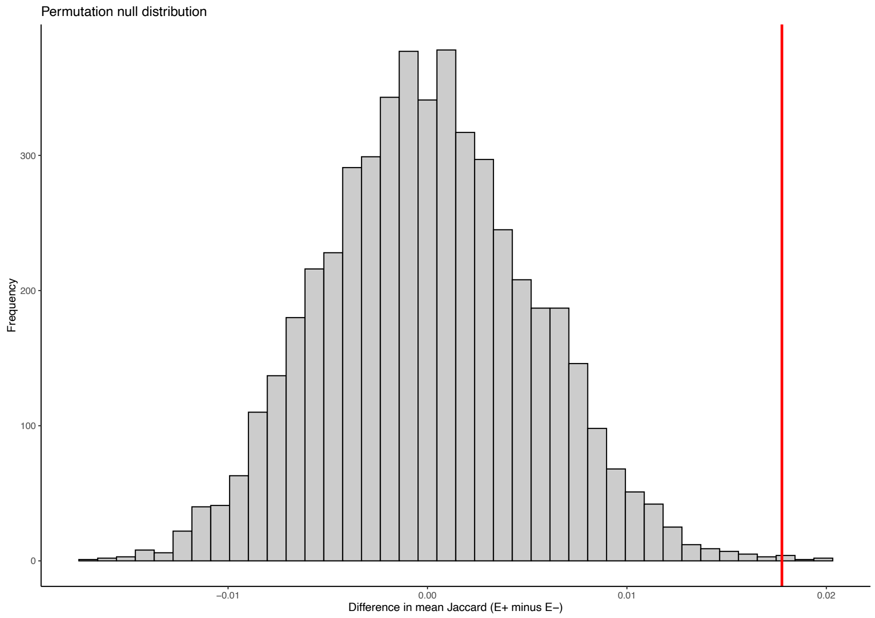


**Figure S13. Permutation test of pairwise niche overlap of *Poa* species**. Pairwise niche overlap was quantified using the Jaccard index calculated across all grid cells of each species range. The histogram shows the null distribution of differences in mean pairwise Jaccard similarity between endophyte-positive (E+) and endophyte-negative (E-) species generated by permuting endophyte labels across species (5,000 permutations). The vertical red line indicates the observed difference in mean Jaccard similarity (E+ minus E-).

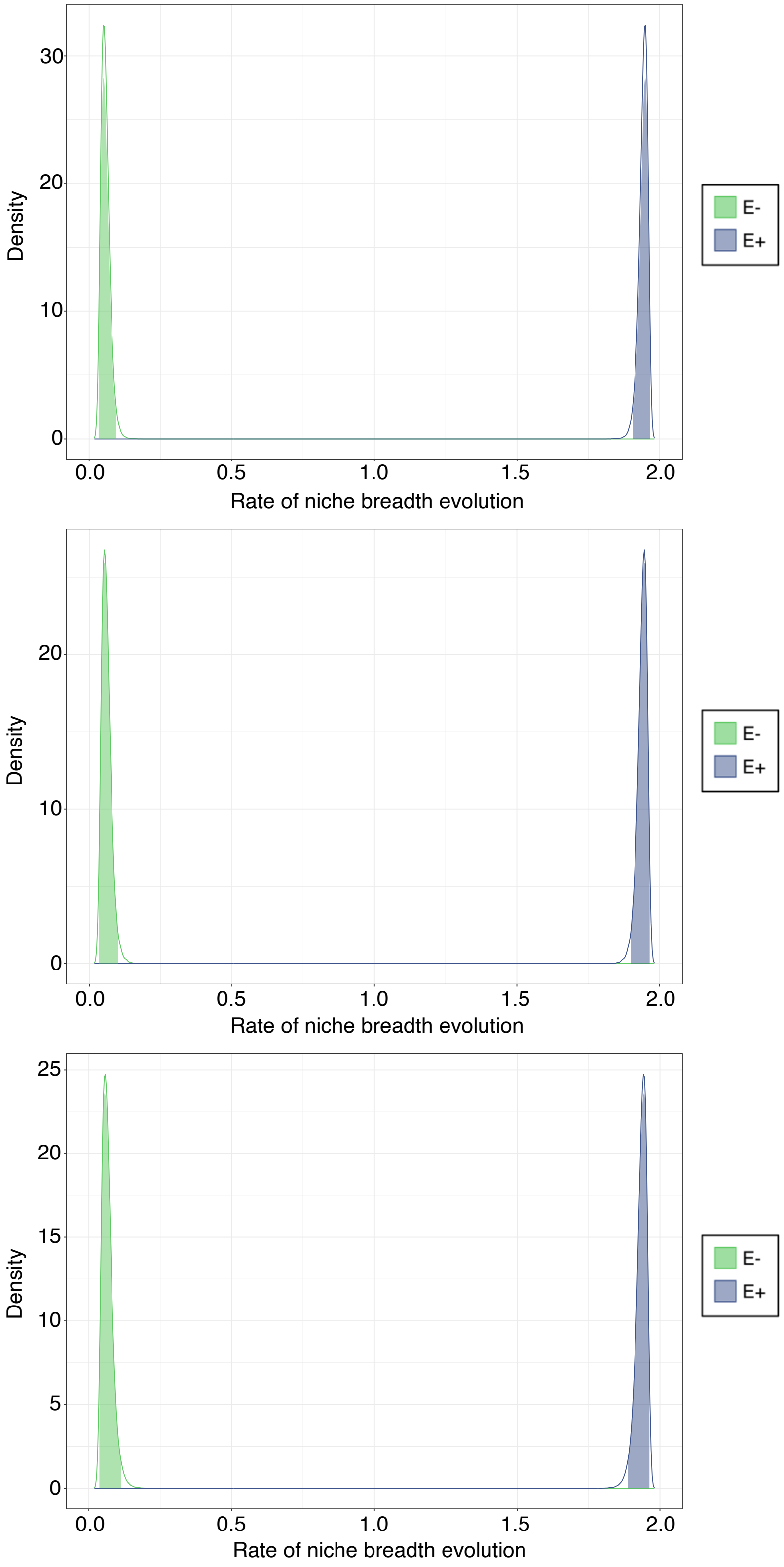


**Figure S14. Estimated rates of niche breadth evolution from MuSSCRat models, testing different priors.** Estimated rates of niche breadth evolution (range size, range on the primary niche axis, and hypervolume) for E- and E+ *Poa* species. Models used different priors on the number of rate shifts of the multivariate continuous response variable, testing (from top to bottom) 5, 10, and 20 shifts.

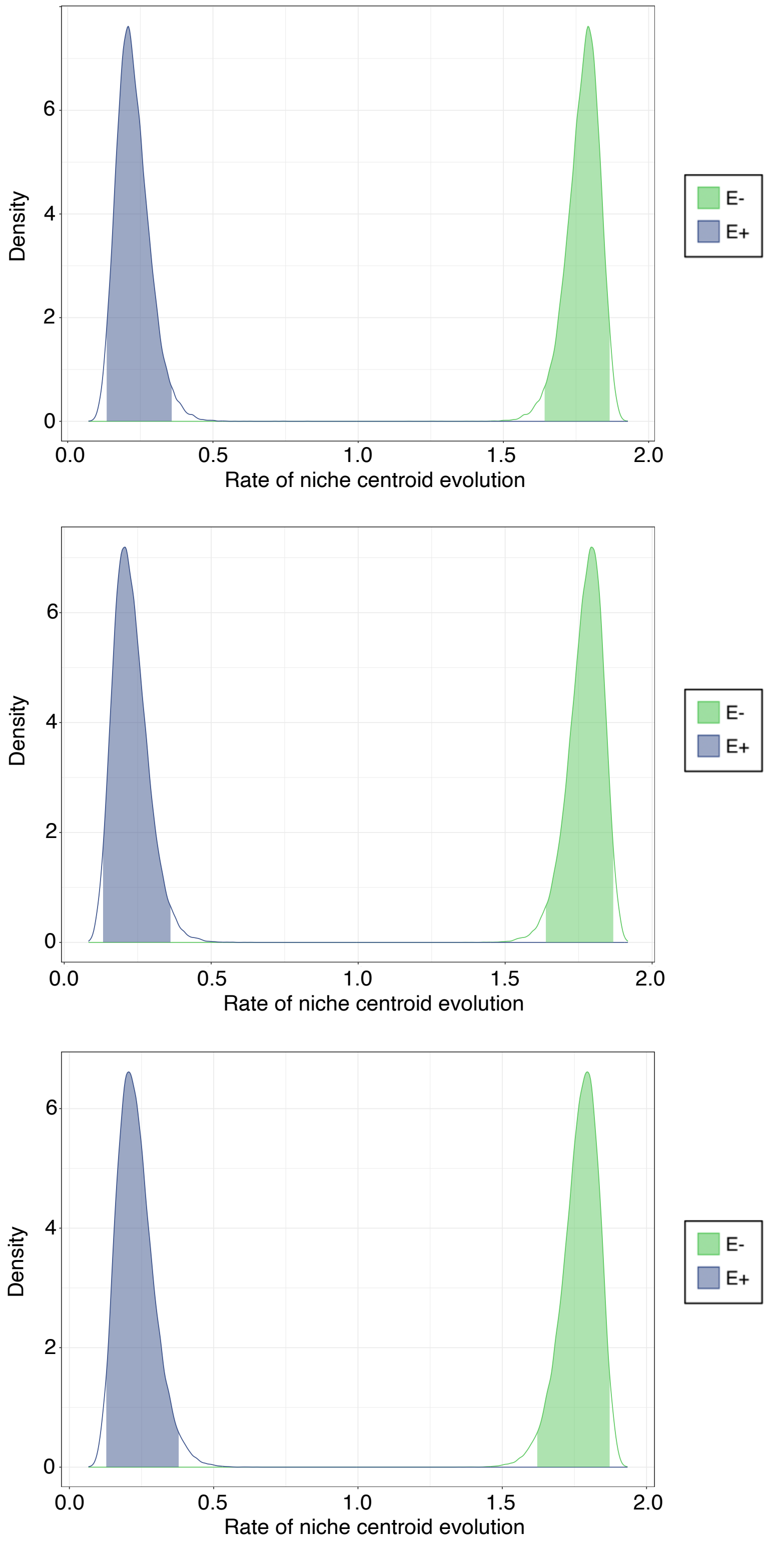


**Figure S15. Estimated rates of niche centroid evolution from MuSSCRat models, testing different priors.** Estimated rates of niche centroid evolution (temperature, precipitation, nitrogen, and clay) for E+ and E- *Poa* species. Models used different priors on the number of rate shifts of the multivariate continuous response variable, testing (from top to bottom) 5, 10, and 20 shifts.

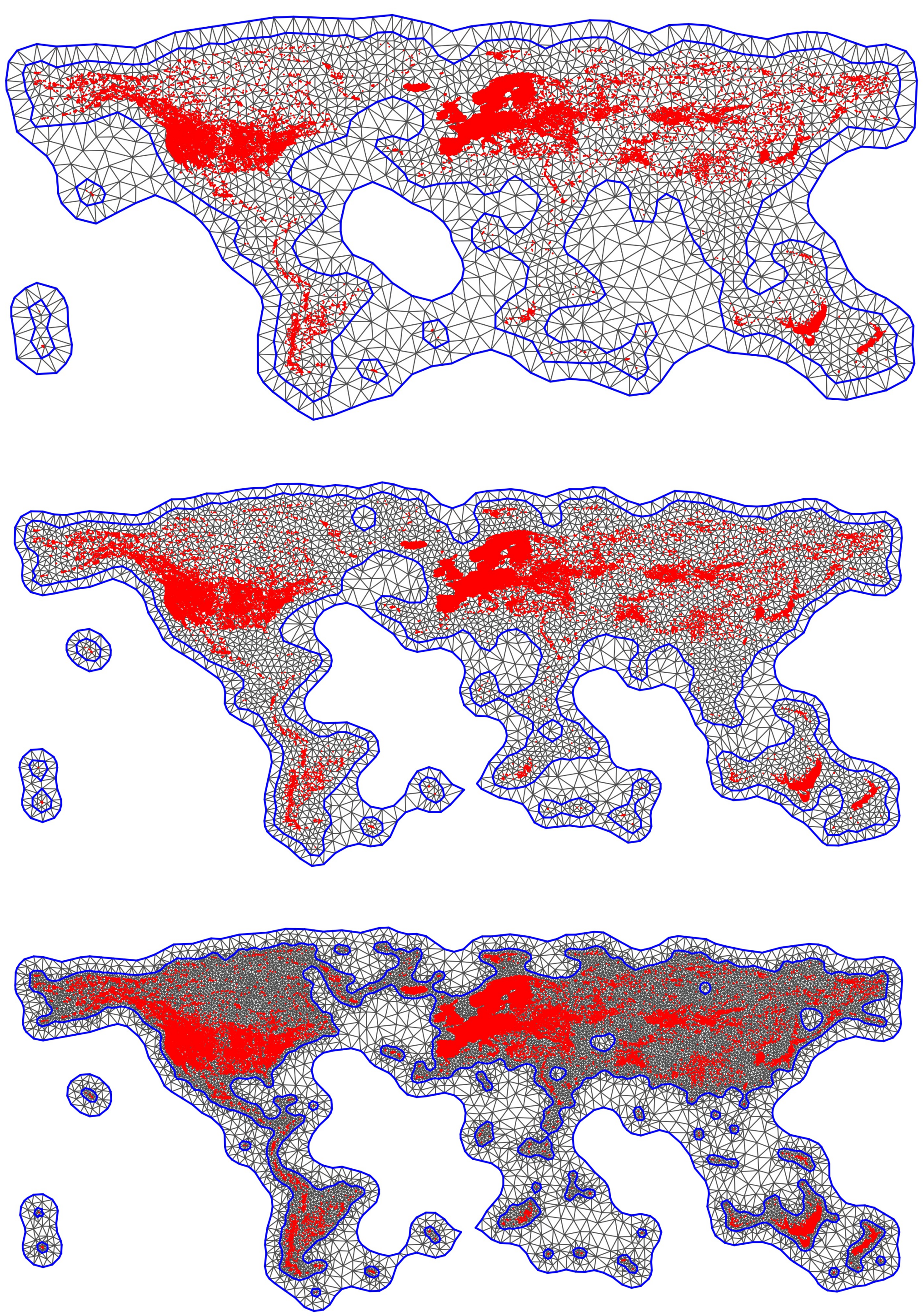

**Figure S16. The three spatial meshes tested for INLA models.** From top to bottom, meshes consisted of 2528, 4688, and 13018 vertices, with the final models using the mesh consisting of 2528 vertices. *Poa* point occurrence records are shown with the red dots.

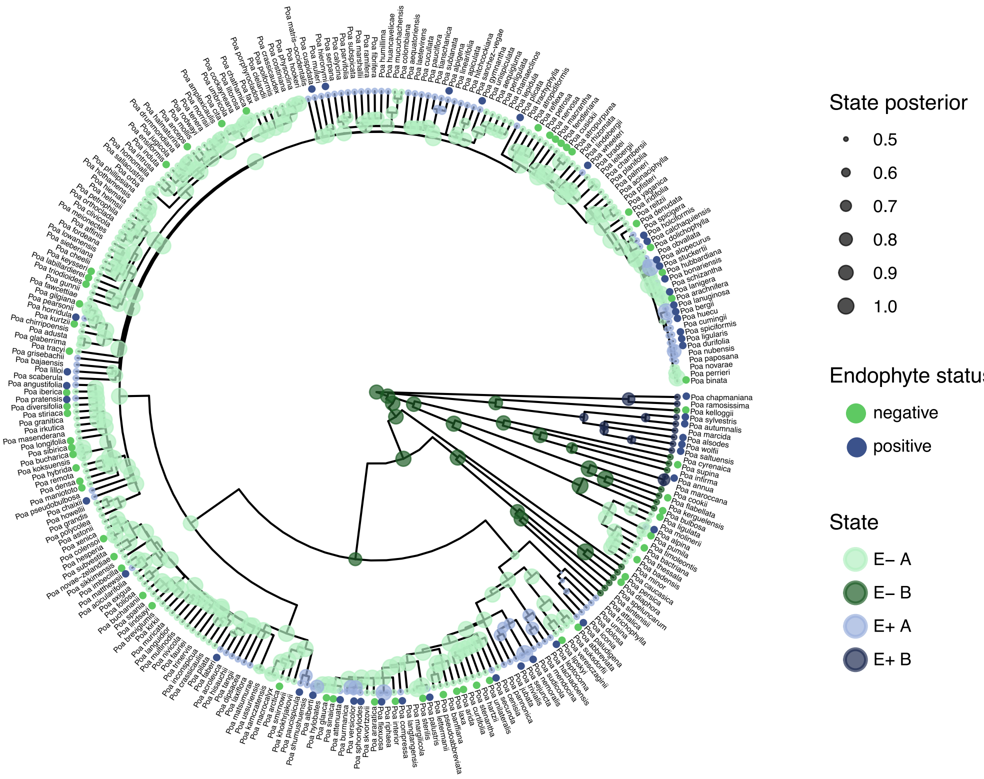


**Figure S17. Ancestral state reconstruction of *Epichloë* endophytes across the *Poa* phylogeny.** The phylogeny displays the ancestral character history of endophytes estimated by the HiSSE model. Here, we used a 280 species tree of *Poa*, and we had endophyte information for 107 of these species (indicated by the opaque green and blue dots at the tips of the tree). States of nodes are indicated by colors of dots over each respective node, with the posterior indicated by the size of the dot.
